# Lung Resident T Cells Across the Spectrum of TB Infection and Disease

**DOI:** 10.64898/2026.08.10.743910

**Authors:** Dylan Kain, GW McElfresh, Katherine H. Rott, Meghan Cansler, Gwendolyn M. Swarbrick, Gerhard Walzl, Nelita Du Plessis, Deborah A. Lewinsohn, Benjamin N. Bimber, David M. Lewinsohn

## Abstract

Tuberculosis (TB) remains the leading infectious cause of death worldwide, yet immunological protection against *Mycobacterium tuberculosis* (Mtb) remain poorly understood. We performed integrated single-cell transcriptomics and functional T-cell cloning on paired bronchoalveolar lavage (BAL) and peripheral blood samples from those exposed to TB, using IGRA and PET-CT to study potentially protective responses across the spectrum of TB infection/disease. Recent Mtb exposure alone was sufficient to remodel the pulmonary T-cell compartment. Longitudinal follow-up demonstrated that all participants who progressed to active TB had baseline PET-CT abnormalities, showing that PET-CT scan can be used to find those at high risk of progression. Furthermore, those who were PET-positive, but did not progress to active TB provide a natural model for protection, and exhibited enrichment of pulmonary cytotoxic CD8-associated T cells, identifying a candidate protective immune program. TAR-seq identified a discrete population of clonally expanded Mtb-responsive T cells and enabled direct linkage of antigen-responsive TCRs to transcriptional state through T cell cloning. Functional T-cell cloning further established a framework connecting antigen specificity, TCR sequence, and pulmonary cell state. Together, these findings provide a comprehensive atlas of human pulmonary T-cell immunity across the spectrum of TB, identify pulmonary immune features associated with durable control of infection, and establish an integrated platform for defining protective T-cell responses to inform next-generation TB vaccine development.

## Introduction

Tuberculosis (TB), caused by *Mycobacterium tuberculosis* (Mtb), remains the leading infectious cause of death worldwide, resulting in 1.23 million deaths in 2024 [1]. Although widespread neonatal vaccination with *Mycobacterium bovis* bacille Calmette–Guérin (BCG) has substantially reduced severe disseminated disease in young children [2], it has not been shown to provide consistent protection against adult pulmonary TB [3], underscoring the urgent need for novel vaccines. Following exposure to Mtb, outcomes vary widely with most individuals successfully containing or possibly eliminate infection, while a minority progress to active TB disease [4]. These divergent outcomes highlight the critical role of the host immune response, and in particular the role of T-cell immunity in protection against TB [5–8]. Although immune responses are most commonly studied in peripheral blood, TB is fundamentally a disease of the lung. The initial interaction between Mtb and the host occurs within the respiratory tract, where immune cells must effectively and specifically recognize and control infection. Increasing evidence suggests that tissue-resident immune populations in the lung provide an effective localized response that may determine infection outcome [9,10]. In this regard, while there has been extensive work characterizing both CD4 and CD8 antigens from peripheral blood [11], there has been no work performed defining the T cell antigens recognized in the lung [12]. Defining these pulmonary T cell responses and the antigens recognized are therefore essential for identifying correlates of protection and informing next-generation TB vaccine design.

Historically, TB has been viewed as a binary state comprising either latent TB infection or active disease. However, advances in microbiological, imaging and immunological techniques have increasingly demonstrated that TB exists along a dynamic spectrum of infection, ranging from complete bacterial clearance through contained infection, microbiologically negative asymptomatic disease, microbiologically positive asymptomatic disease, and symptomatic disease [13]. Specifically, asymptomatic TB has emerged as a far more prevalent stage of disease than previously appreciated and may occur in individuals with either microbiologically confirmed disease or radiographic evidence of disease despite negative microbiological testing [14–16]. This spectrum likely reflects both the degree of exposure and the diverse immune outcomes that occur following Mtb exposure rather than discrete biological states. Understanding immune responses during asymptomatic infection is essential because these individuals represent a naturally occurring model of successful immune control despite ongoing pulmonary infection and inflammation. Furthermore, epidemiological and modeling studies suggest that a substantial proportion of Mtb transmission may arise from individuals with asymptomatic, microbiologically positive disease [17,18]. Moreover, preventing progression through this stage may be necessary for vaccines intended to interrupt transmission. Current clinical tools—including interferon-γ release assays (IGRAs) and the tuberculin skin test—identify immunological sensitization to Mtb but do not provide insight into ongoing Mtb infection or accurately predict who will progress to developed TB disease. Consequently, new approaches are required to investigate the biological heterogeneity underlying these diverse clinical outcomes.

One of the most powerful approaches for studying early human TB has been the integration of longitudinal household contact cohorts with positron emission tomography-computed tomography (PET-CT) imaging. PET-CT can detect metabolically active pulmonary lesions long before the development of symptoms or microbiological confirmation, providing an opportunity to identify individuals with early, localized pulmonary inflammation following recent Mtb exposure. Importantly, longitudinal studies of household contacts have demonstrated that PET-CT abnormalities are predictive of subsequent progression to microbiologically confirmed TB, while the majority of individuals with PET-positive lesions ultimately control infection without developing active disease [19]. This unique population therefore offers an exceptional opportunity to investigate pulmonary T cell responses, inclusive of their phenotype and antigen specificity, associated with successful containment of Mtb during the earliest stages of infection. By coupling PET-CT imaging with bronchoalveolar lavage, peripheral blood sampling and longitudinal clinical follow-up, it becomes possible to characterize these T cell responses directly at the site of infection and distinguish mechanisms associated with protection from those preceding disease progression, as well as identifying the antigens recognized in this protective context. Such studies may ultimately inform the rational development of vaccines capable of preventing pulmonary TB.

To that end, we conducted a household contact (HHC) study in Cape Town, South Africa. Combined single-cell gene expression (scGEX-seq), TCR sequencing (sc-TCR-seq) and CITE-seq was performed on peripheral blood mononuclear cell (PBMC) samples and matched non-adherent BAL samples from individuals with active TB, exposed household contacts (HHCs) and community controls (CC). HHCs were further stratified by serial IGRA and PET-CT imaging to distinguish individuals with IGRA conversion and subclinical lung inflammation (PET-CT-positive). Participants were followed for three years to determine progression to active TB disease. We identified marked transcriptional and TCR differences between circulating and lung-resident T cells and demonstrated that lung-resident T-cell states varied across the spectrum of TB infection and disease. Compared with other groups, community controls exhibited increased expression of naïve and stem-like T-cell markers, suggesting recent Mtb exposure is sufficient to reshape the pulmonary T-cell compartment. Progression to active TB occurred exclusively in individuals with baseline PET-CT positivity, and comparing these progressors to PET-positive non-progressors, the non-progressors had an enriched cytotoxic CD8 T cell cluster associated with protection against TB. Using TAR-seq, a method that identifies antigen-responsive T cells through paired TCR sequencing and activation profiling [20], we found that only a small proportion of BAL T cells responded to ex vivo Mtb stimulation, although these responses were enriched in active TB and PET-CT-positive individuals. Finally, by generating T-cell clones from BAL samples of individuals with active TB, we isolated Mtb-specific T cells, identified their cognate antigens, and demonstrated that these clonotypes could be tracked within the single-cell datasets following antigen stimulation. Overall, this work provides a comprehensive single-cell atlas of human lung-resident T cells across the spectrum of Mtb exposure, infection, asymptomatic disease, and active TB. These findings reveal how the pulmonary T-cell compartment is remodeled during early infection, identify immune features associated with progression risk and potential protection, and establish a framework for tracking antigen-specific T-cell responses directly within the human lung.

## Results

### Study design and characterization of household contacts across the spectrum of tuberculosis infection

To investigate pulmonary T-cell responses across the spectrum of Mtb exposure and disease, we enrolled adults from Cape Town, South Africa with newly diagnosed active TB (n=10), their asymptomatic and microbiologically negative HHCs (n=32), and IGRA-positive community controls (CC) (n=11) without known recent exposure (Figure 1a) [21]. HHCs were further stratified using serial IGRA testing (months 0 and 3) and PET-CT into the following three groups; IGRA-positive at month 0 and 3 with negative PET-CT scan (PET^NEG^; n=11), IGRA conversion with negative PET-CT scan (Converter; n=8), and those with a positive PET-CT scan (PET^POS^; n=13). Participant characteristics are summarized in Table 1 with active TB participants having significantly lower BMI. To study lung-resident populations, matched PBMC and bronchoalveolar lavage (BAL) samples were collected from all participants, and BAL cell differentials at time of collection did not differ significantly between groups (Supplemental Figure 1a,b, Supplemental Table 1). To enrich for lymphocytes, BAL samples underwent macrophage depletion by adherence followed by flow cytometric sorting of BAL and PBMC cells based on size (Supplemental Figure 1c). To identify Mtb-responsive T cells, enriched lymphocytes were then exposure to autologous dendritic cells which were either uninfected (“unstimulated”) or infected overnight with an Mtb auxotroph strain (“stimulated”) [22]. Samples then had combined single-cell gene expression (scGEX-seq), TCR-sequencing (scTCR-seq) and cell surface protein expression via CITE-seq.

**Figure 1:**
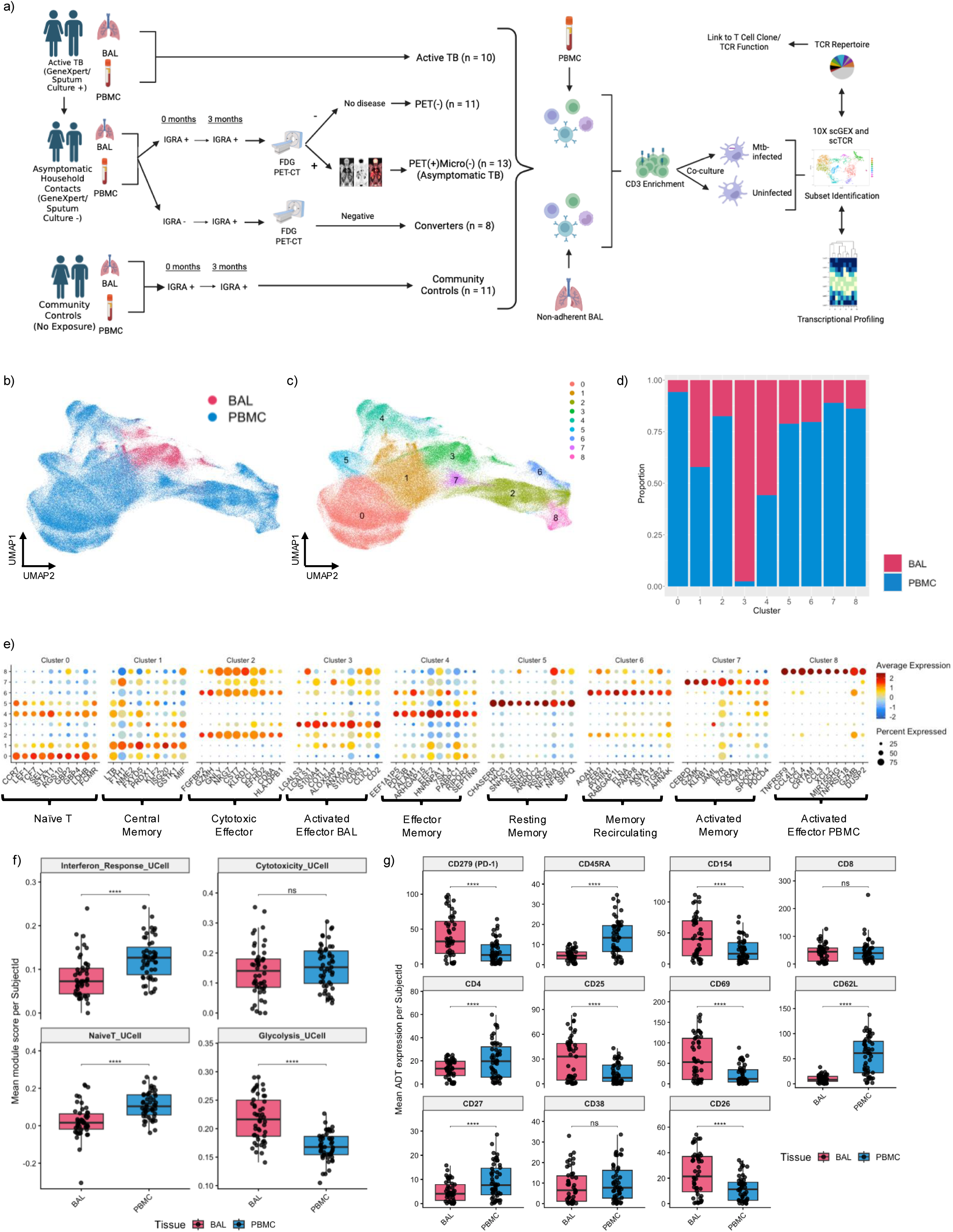
BAL and PBMC T Cells Have Distinct Transcriptional Profiles. a) Study design and experimental workflow. Adults with microbiologically confirmed active TB, household contacts (HHCs), and IGRA-positive community controls (CCs) were enrolled. HHCs underwent serial IGRA testing and PET-CT imaging and were classified as PET-CT negative (PET^NEG^), PET-CT positive (PET^POS^), or IGRA converters (who also had negative PET-CT scan). BAL and PBMC samples were collected from all participants. Following macrophage depletion and lymphocyte enrichment, cells were co-cultured with autologous dendritic cells that were either uninfected (unstimulated) or infected overnight with an Mtb auxotroph strain (stimulated). Samples were analyzed using single-cell gene expression (scGEX-seq), paired TCR sequencing (scTCR-seq) and CITE-seq. b) UMAP reduction of BAL and PBMC derived T cells from all participants (n=53). c) Unbiased Leiden clustering of BAL and PBMC derived T cells. d) Proportion of each cluster derived from BAL or PBMC samples. e) Dotplot of key marker genes derived from DEG analysis and established lineage/phenotypic markers. Color represents average expression, and size represents percent of T cells expressing that gene. f) Interferon Response (*IFI6, IFI27, MX1, ISG15, STAT1, MX2, IFIT3*), Cytotoxicity (*PRF1, GNLY, NKG7, GZMA, GZMB, GZMH, GZMK* and *GZMM*), Naïve T (*CTSG, CA6, GSTT1, LEF1, RGS10* and *TMIGD2*) and Glycolysis (*ALDOA, BPGM, ENO1, ENO2, GAPDH, HK1, HK2, HKDC1, PFKL, PGAM1, PGAM2, PGK1, PKLR, PKM, TPI1*) gene module scores for each participant segregated based on T cell compartment origin. Each point represents an individual participant. Boxes indicate the median and interquartile range. P values were calculated using the Wilcoxon rank-sum test with Benjamini–Hochberg (BH) correction for multiple testing. *p-value < 0.05; **p-value < 0.01; ***p-value < 0.001; ****p-value < 0.0001. Interferon Response p = 5.09e^-5^, Cytotoxicity p = 0.216, Naïve T p = 9.79e^-6^, Glycolysis p = 7.84 e^-8^. g) CITE-seq surface protein expression in BAL and PBMC T cells. Each point represents an individual participant. Boxes indicate the median and interquartile range. P values were calculated using the Wilcoxon rank-sum test with Benjamini–Hochberg (BH) correction for multiple testing. *p-value < 0.05; **p-value < 0.01; ***p-value < 0.001; ****p-value < 0.0001. CD279 (PD-1) p = 5.76e^-7^, CD45RA p = 4.59e^-7^, CD154 p = 2.51e^-6^, CD8 p = 0.571, CD4 p = 6.32e^-5^, CD25 p = 7.24e^-7^, CD69 p = 2.28e^-7^, CD62L p = 5.51e^-8^, CD27 p = 1.89e^-5^, CD38 p = 0.57, CD26 p = 7.75e^-7^.

**Table 1:**
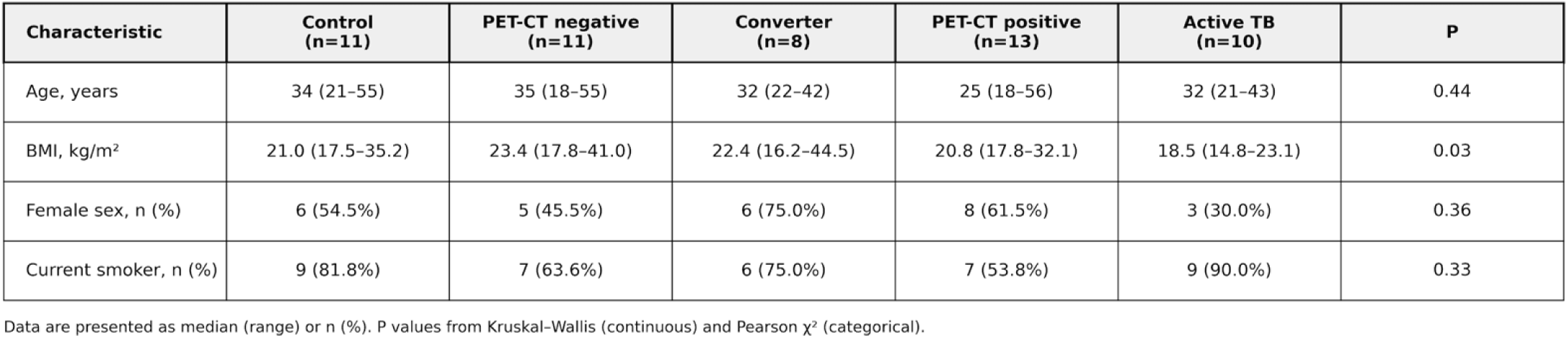
Participant Demographics by Donor Group. Data are presented as median (IQR) or *n* (%). P values were calculated using the Kruskal–Wallis test for continuous variables and Fisher’s exact test for categorical variables.

### BAL and PBMC T Cells Have Unique Gene Expression Profiles

After quality control steps there were 37,952 BAL-derived and 297,557 PBMC-derived T cells included in the complete analysis. Individual cell counts are shown in Supplemental Figure 2a. To define tissue-resident program, unstimulated BAL (20,096 cells) and PBMC (148,730 cells) T cells were first examined. Unbiased Leiden clustering identified nine transcriptional clusters (Figure 1b–d, Supplemental Figure 2b), with most BAL T cells localizing to cluster 3. This cluster preferentially expessed genes associated with activation and tissue adaptation, including *LGALS3*, *S100A4*, and *COTL1* (Figure 1e, Supplemental Table 2) [23–25]. Pseudobulk differential expression analysis confirmed distinct compartment-specific transcriptional programs (Supplemental Figure 2c, Supplemental Table 3). BAL T cells preferentially expressed activation and tissue-resident genes (*LGALS3, S100A4, COTL1*), whereas PBMC T cells were enriched for interferon-stimulated (*IFI6, IFI27, ISG15*) and circulating/naïve-associated genes (*CCR7, LEF1, IL7R*). Gene module analysis demonstrated lower interferon-response scores and higher glycolysis and naïve T-cell scores in BAL T cells (Figure 1f) [26]. CITE-seq further distinguished BAL T cells by increased expression of CD69, CD103, HLA-DR, PD-1, and CD8, whereas PBMC T cells preferentially expressed CD45RA, CD62L, CD27, and CD4 (Figure 1g). To determine whether these differences reflected compartment origin rather than disease state, BAL and PBMC T cells from community controls were analyzed separately (Supplemental Figure 3, Supplemental Table 4). The compartment-specific transcriptional programs were preserved, with BAL T cells remaining enriched for tissue-resident and effector genes (*CXCR6, GZMK, XCL1, KLRC1*) and PBMC T cells for interferon-response and circulating T-cell genes (*IFI6, IFI27, IFIT2, GBP4, STAT1, MAL, TRABD2A*). Overall, these findings demonstrate that lung-resident T cells are phenotypically distinct from circulating T cells, exhibiting features of tissue residency and sustained activation [23,27,28].

### BAL T Cells Have Unique Gene Expression Profiles Across the Spectrum of Mtb Infection/Disease

To explore lung-resident T cells, BAL cells were reclustered to define transcriptional changes across the spectrum of *Mtb* exposure and disease (Figure 2a). Nine BAL T-cell clusters were identified (Figure 2b) with varied proportion of each group (Figure 2c). Key marker genes derived from DEG analysis and established lineage/phenotypic markers are shown in Figure 2d (Supplemental Table 5). Community controls were enriched for naïve/central memory (cluster 5; *CCR7, TCF7, LEF1*) and terminal cytotoxic (cluster 7; *PRF1*) populations, PET^NEG^ participants for an activated Treg cluster (cluster 8; *FOXP3, IL2RA, CTLA4*), converters for an interferon-stimulated cluster (cluster 4; *IFIT1, ISG15, MX1*), PET^POS^ participants for a type 2-like memory cluster (cluster 6; *GATA3, IL17RB, HPGDS*), and active TB for an activated memory cluster (cluster 2). Pseudobulk differential expression demonstrated distinct transcriptional programs at each stage of *Mtb* exposure and disease, with most DEGs unique to individual comparisons (Figure 2e, Supplemental Table 6). Gene ontology (GO) analysis identified group-specific biological pathways, with progression from leukocyte migration and homeostasis in CC and PET^NEG^ participants toward inflammatory and interferon-associated pathways in PET^POS^ and innate inflammatory signaling in active TB (Figure 2f, Supplemental Table 7). Together, these findings demonstrate progressive inflammatory remodeling of pulmonary T-cell transcriptional programs across the spectrum of *Mtb* exposure and disease.

**Figure 2:**
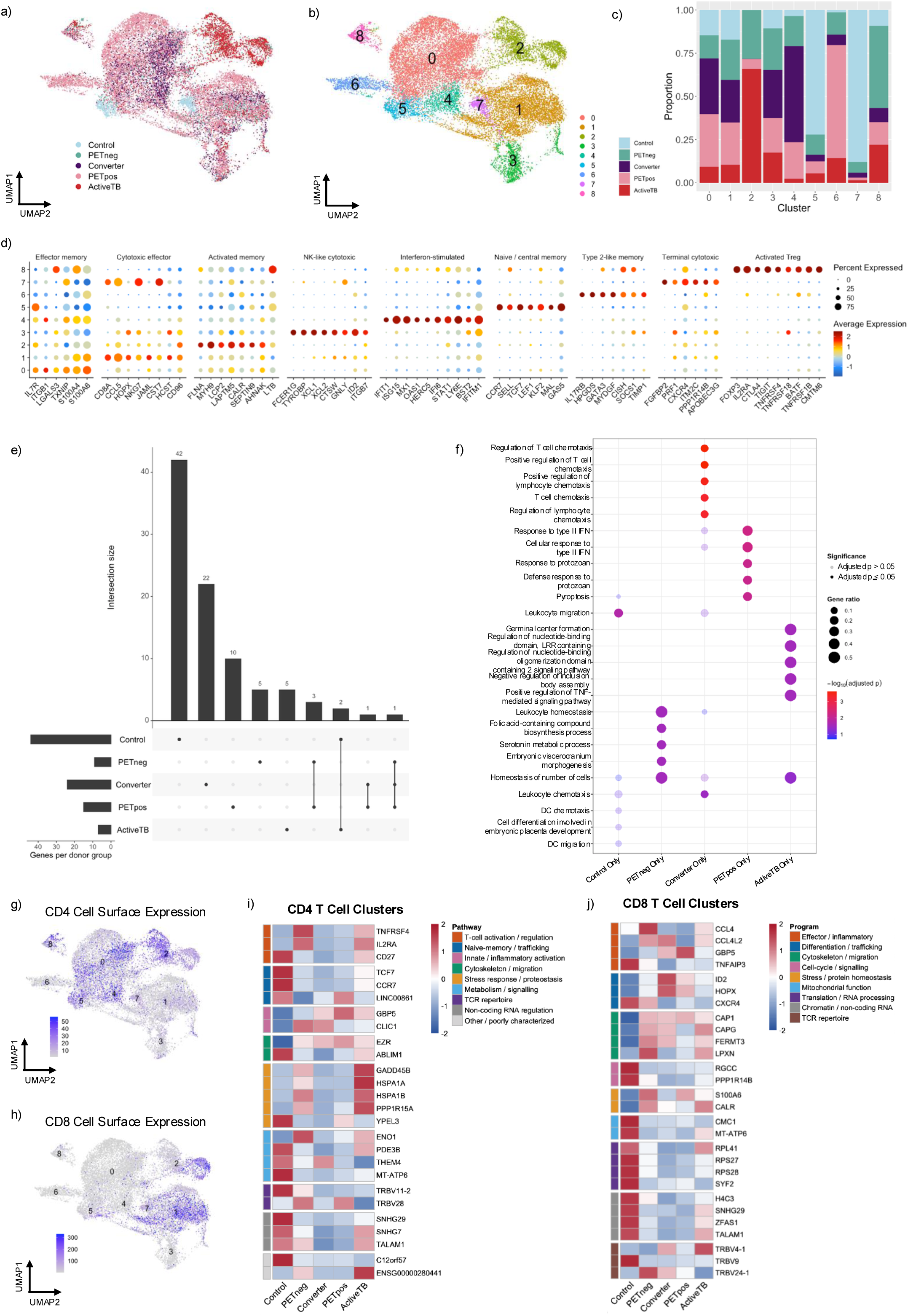
BAL T Cells Have Distinct Transcriptional Profiles Across the Spectrum of TB Infection/Disease. a) UMAP of re-clustered BAL T cells from each participant group. b) Unbiased Leiden clustering of BAL T cells. c) Proportion of each cluster derived from individual participant groups. d) Dotplot of key marker genes derived from DEG analysis and established lineage/phenotypic markers. Color represents average expression, and size represents percent of T cells expressing that gene. e) UpSet plot showing the overlap of significantly differentially expressed genes identified by pseudobulk differential expression analysis across pairwise donor group comparisons. Bars indicate the number of unique or shared differentially expressed genes, with connected dots indicating the comparisons contributing to each intersection. Differential expression analysis was performed on pseudobulk profiles using DESeq2, with significance defined as Benjamini–Hochberg-adjusted *P* < 0.05. f) Gene Ontology (GO) biological process enrichment analysis of pseudobulk differentially expressed genes across donor group comparisons. Dot size represents the number of differentially expressed genes associated with each pathway, and color indicates the adjusted *P* value. GO enrichment was performed using Benjamini–Hochberg correction for multiple testing, with significantly enriched pathways defined as adjusted *P* < 0.05. g) UMAP of BAL T cells coloured by CD4 CITE-seq expression. h) UMAP of BAL T cells colored by CD8 CITE-seq expression. i) Heatmap of significantly differentially expressed genes identified by pseudobulk differential expression analysis of CD4 T cells clusters (cluster 0, 4, 5, 8) across donor groups (DESeq2, BH-adjusted *P* < 0.05). j) Heatmap of significantly differentially expressed genes identified by pseudobulk differential expression analysis of CD8 T cells clusters (cluster 1, 7) across donor groups (DESeq2, BH-adjusted *P* < 0.05).

### Pulmonary CD4 and CD8 T cells exhibit distinct transcriptional profile across the spectrum of TB infection/disease

CD4 and CD8 T cells often have distinct responses and so to explore them separately, surface protein expression identified clusters 0, 4, 5, and 8 as predominantly CD4 T cells, clusters 1 and 7 as CD8 T cells, cluster 2 as mixed CD4/CD8, and clusters 3 and 6 as double-negative (Figure 2g,h; Supplemental Figure 4a). Pseudobulk analysis demonstrated distinct transcriptional changes within both CD4 and CD8 compartments (Supplemental Figures 4–5; Supplemental Tables 8–9). Within CD4 T cell clusters, community controls preferentially expressed naïve and lymphoid-trafficking genes (*TCF7, CCR7*), whereas PET^NEG^ upregulated activation/regulatory genes (*TNFRSF4, IL2RA*), converters inflammatory genes (*GBP5, EZR*), PET^POS^ metabolic genes (*ENO1, PDE3B*), and active TB stress-response genes (*GADD45B, HSPA1A/B*). CD8 T cells also underwent progressive changes, with PET^NEG^ enriched for effector genes (*CCL4*), converters for differentiation-associated genes (*GBP5, HOPX, ID2*), PET^POS^ for stress-response genes (*S100A6, CALR*), and community controls for genes associated with mitochondrial function and protein translation. GO analysis demonstrated enrichment of activation, inflammatory, metabolic, and TCR-signaling pathways within CD4 T cells, whereas CD8 T cells preferentially upregulated pathways related to effector function, differentiation, cytoskeletal remodeling, and TCR repertoire (Figure 2g,h).

### HHC Exposure to TB is Sufficient to Reshape the Pulmonary T-cell Compartment

Because community controls were enriched for a naïve/stem-like T-cell cluster, we next examined canonical naïve-associated genes across participants (Figure 3a). CC exhibited significantly higher naïve/stem-like module scores than converters (p=0.0186) and active TB participants (p=0.0403), with similar trends in PET^NEG^ and PET^POS^ participants (Figure 3b). CITE-seq confirmed increased frequencies of CCR7 CD45RA naïve-like and CCR7 CD45RA central memory-like cells in community controls (Figure 3c–e) [29]. Cell-type annotation [30] similarly demonstrated enrichment of naïve CD4 T cells (p=0.041) and a similar trend in naïve CD8 T cells (Figure 3f). These findings indicate that pulmonary T-cell differentiation begins following *Mtb* exposure, even before PET-CT abnormalities or IGRA conversion.

**Figure 3.**
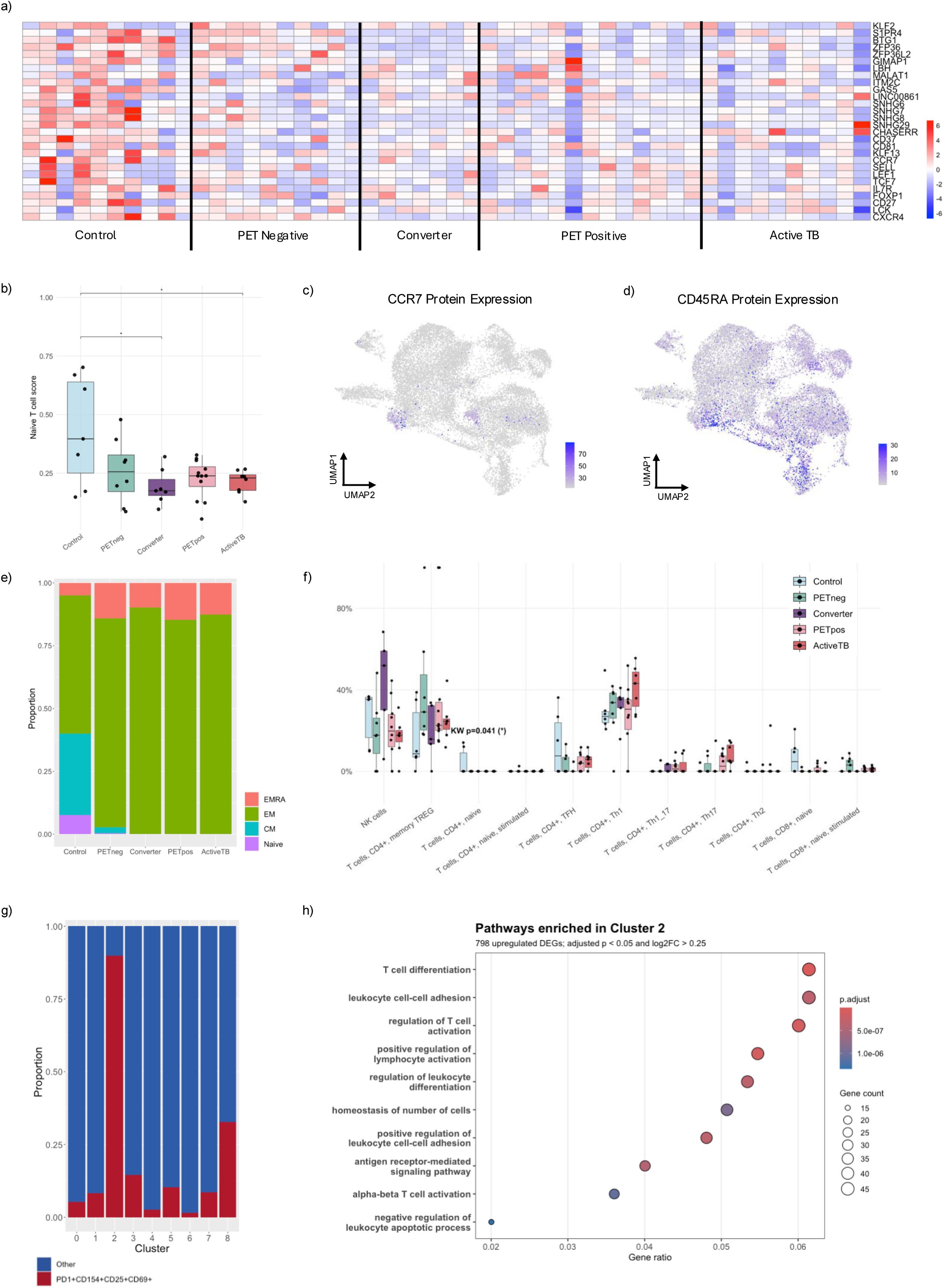
HHC Exposure is Sufficient to Reshape the Pulmonary T Cell Compartment. a) Heatmap of cannoncial naïve and stem-like T cell genes with each column representing individual participants. b) Quantification of naïve and stem-like genes from heatmap above combined into a gene module score, with each dot representing an individual participant. Boxes indicate the median and interquartile range. P values were calculated using the Kruskal-Wallis test. *p-value < 0.05. Control vs converter p = 0.0186, control vs active TB p = 0.040 c) UMAP of BAL T cells coloured by CCR7 CITE-seq expression. d) UMAP of BAL T cells colored by CD45RA CITE-seq expression. e) Proportion of BAL T cells classified into naïve (CCR7 CD45RA), central memory (CCR7 CD45RA), effector memory (EM; CCR7 CD45RA), and terminally differentiated effector memory (EMRA; CCR7 CD45RA) subsets based on CITE-seq expression of CCR7 and CD45RA. f) Proportion of BAL T cells assigned to reference T-cell subsets using the CellTypist transfer algorithm [30] across donor groups. P values were calculated using the Kruskal–Wallis test. Only significant comparisons are indicated (*P* < 0.05) g) Proportion of cells within each BAL T-cell cluster co-expressing PD-1, CD154, CD25, and CD69 as measured by CITE-seq. Bars represent the fraction of cells positive for all four activation markers within each cluster. h) Gene Ontology (GO) biological process enrichment analysis of Cluster 2 marker genes identified by differential expression analysis (Wilcoxon rank-sum test, BH-adjusted p < 0.05, log_2_FC > 0.25). Dot size represents gene count, and color indicates the BH-adjusted p value.

### Active TB is associated with an activated pulmonary T-cell population

Cluster 2, enriched in active TB, comprised both CD4 and CD8 T cells (Supplemental Figure 4a–c) and expressed high levels of PD-1, CD154, CD25, and CD69 (Figure 3g), consistent with recent antigen-driven activation and pulmonary tissue retention [31–33]. Gene ontology analysis demonstrated enrichment of T-cell activation, antigen receptor signaling, leukocyte differentiation, and survival pathways (Figure 3h), identifying this cluster as a highly activated pulmonary T-cell population that expands during active TB.

### The pulmonary T-cell repertoire is largely distinct from the peripheral circulation and is less diverse

Comparison of paired BAL and PBMC TCR repertoires demonstrated that only 5–10% of unique BAL clonotypes were also detected in matched blood, with no differences across participant groups (Figure 4a). Among shared clonotypes, relative frequencies correlated only weakly between compartments (Spearman’s ρ = –0.18-0.28), with the strongest correlation observed in CCs (ρ=0.28, p=0.012) (Figure 4b). The weaker correlations observed in HHC and active TB may suggest increasing expansion of tissue-specific clonotypes following Mtb exposure and infection, although this will require confirmation in larger cohorts. Inverse Simpson diversity did not differ significantly between participant groups in either BAL or PBMC (Figure 4c).

**Figure 4.**
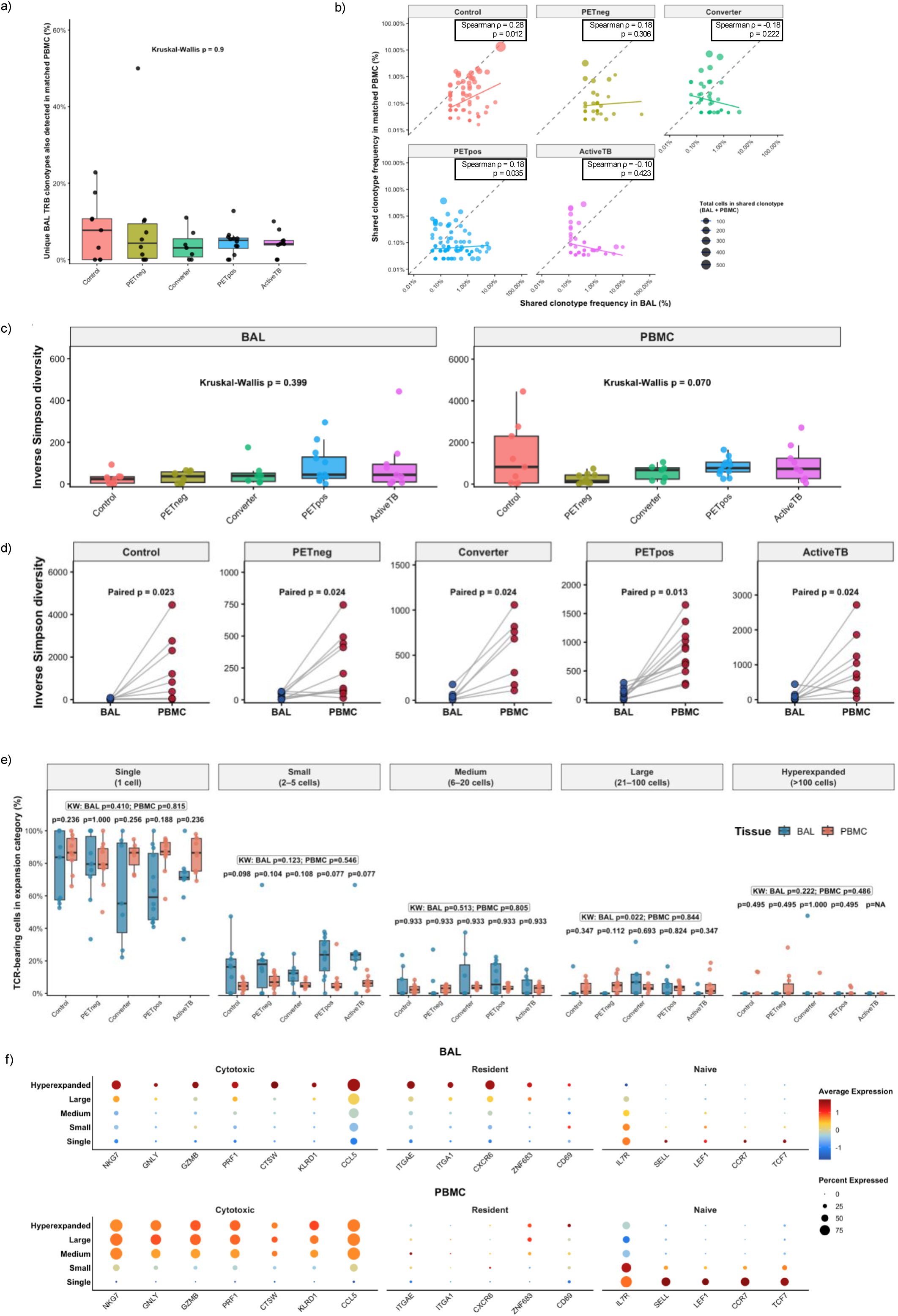
BAL and PBMC T Cells have Distinct TCR Repetoires. a) Percentage of unique BAL TRB clonotypes also detected in matched PBMC from the same participant across donor groups. Clonotypes were defined by identical TRB CDR3 sequences in cells containing at least one TRA chain and exactly one TRB chain. Each point represents an individual participant; boxes indicate the median and interquartile range. P values were calculated using the Kruskal–Wallis test. b) Frequency of shared TRB clonotypes in matched BAL and PBMC samples across donor groups. Each point represents a shared TRB clonotype, with point size proportional to the total number of cells carrying that clonotype across both tissues. Clonotypes were defined by identical TRB CDR3 sequences in cells containing at least one TRA chain and exactly one TRB chain. Frequencies were calculated as the percentage of T cells within each tissue belonging to the shared clonotype. Spearman’s rank correlation coefficient (ρ) and corresponding *P* values are shown for each donor group. c) Inverse Simpson diversity of the TRB repertoire in BAL and PBMC across donor groups. Each point represents an individual participant; boxes indicate the median and interquartile range. P values were calculated using the Kruskal–Wallis test, with pairwise comparisons performed using Dunn’s test with Benjamini–Hochberg (BH) correction. d) Paired comparison of TRB repertoire Inverse Simpson diversity between matched BAL and PBMC samples within each donor group. Each point represents an individual participant, with paired samples connected by lines. P values were calculated using the paired Wilcoxon signed-rank test with BH correction. e) Proportion of TCR-bearing cells within each clonal expansion category in matched BAL and PBMC samples across donor groups. Clonotypes were classified as single (1 cell), small (2–5 cells), medium (6–20 cells), large (21–100 cells), or hyperexpanded (>100 cells). Each point represents an individual participant; boxes indicate the median and interquartile range. Donor-group differences within each tissue were assessed using the Kruskal–Wallis test, with Dunn’s test and BH correction for pairwise comparisons. BAL and PBMC were compared within donor groups using paired Wilcoxon signed-rank tests with BH correction. f) Dot plots showing expression of cytotoxic, tissue-residency, and naïve T-cell-associated genes across TCR clonal expansion categories in BAL and PBMC. Dot size represents the percentage of cells expressing each gene, and color indicates scaled average expression. Clonotypes were categorized as single (1 cell), small (2–5 cells), medium (6–20 cells), large (21–100 cells), or hyperexpanded (>100 cells).

However, paired analysis demonstrated consistently greater TCR diversity in PBMC than BAL across all participant groups (Figure 4d), indicating that the pulmonary T-cell compartment contains a less diverse repertoire consistent with selective recruitment and local expansion of antigen-experienced clonotypes.

### Expanded pulmonary T-cell clonotypes acquire cytotoxic and tissue-resident phenotypes

To determine whether T-cell phenotype varied with clonal expansion, clonotypes were classified as singleton, small (2–5 cells), medium (6–20 cells), large (21–100 cells), or hyperexpanded (>100 cells) [34]. Singleton clonotypes predominated in both BAL and PBMC, although PETPOS and active TB participants showed a trend toward more expanded BAL clonotypes (Figure 4e). Increasing clonal expansion was associated with higher expression of cytotoxic genes (*NKG7, GNLY, GZMB, PRF1*) and lower expression of naïve-associated genes (*IL7R, CCR7, TCF7*) in both compartments (Figure 4f). In contrast, tissue-resident markers (*ITGAE, CXCR6, ZNF683*) increased with clonal expansion only in BAL, indicating acquisition of a tissue-resident program during pulmonary clonal expansion.

### Mtb stimulation induces a discrete effector transcriptional program in BAL T cells that can be identified by TAR-seq

To define transcriptional programs induced by Mtb stimulation, unstimulated and Mtb-stimulated BAL T cells were integrated and reclustered. Although stimulated and unstimulated cells largely overlapped within the UMAP and all participant groups contributed to each cluster (Figure 5a–c), cluster 8 consisted almost exclusively of stimulated cells (Figure 5d), indicating that only a subset of pulmonary T cells mounted a robust response to Mtb stimulation. Cluster annotation identified diverse pulmonary T-cell states (Figure 5e, Supplemental Table 10).

**Figure 5.**
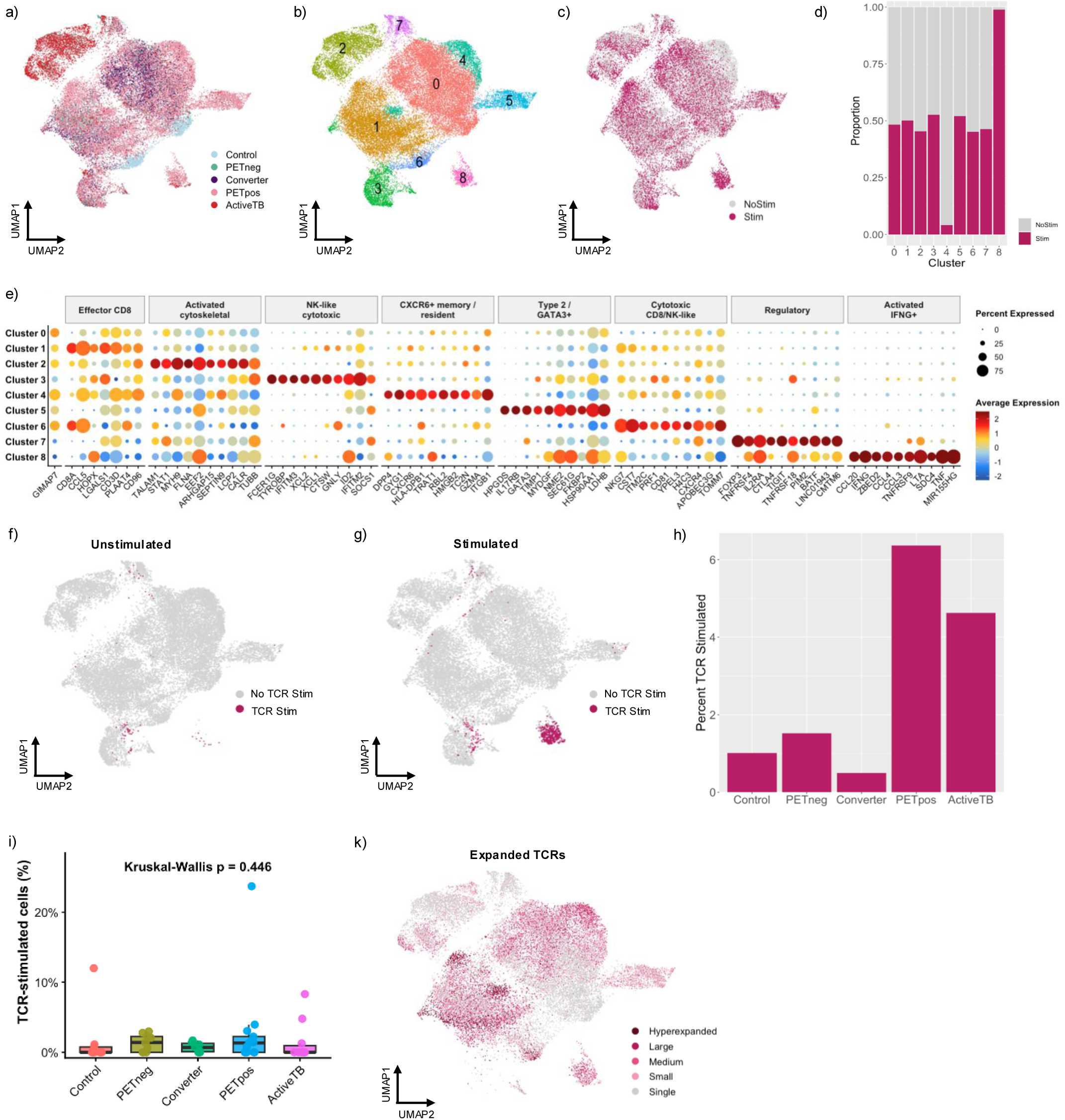
Mtb stimulation identifies a discrete clonally expanded, TCR-responsive pulmonary T-cell population. a) UMAP of integrated unstimulated and Mtb-stimulated BAL T cells colored by donor group. b) UMAP of integrated BAL T cells colored by unbiased Leiden clusters. c) UMAP of integrated BAL T cells colored by stimulation status (unstimulated or Mtb stimulated). d) Proportion of cells within each cluster derived from unstimulated or Mtb-stimulated BAL samples. e) Dot plot showing expression of marker genes used to annotate BAL T-cell clusters. Dot size represents the percentage of cells expressing each gene and color indicates scaled average expression. f) UMAP of unstimulated BAL T cells colored by TAR-seq classification [20] (TCR stimulated or not TCR stimulated). g) UMAP of Mtb-stimulated BAL T cells colored by TAR-seq classification. h) Percentage of Mtb-stimulated BAL T cells classified as TCR stimulated by TAR-seq across donor groups. i) Percentage of Mtb-stimulated BAL T cells classified as TCR stimulated for each participant. Each point represents an individual participant; boxes indicate the median and interquartile range. P values were calculated using the Kruskal–Wallis test, with pairwise comparisons performed using Dunn’s test with Benjamini–Hochberg (BH) correction. j) UMAP showing the distribution of cells lacking a complete paired TCR and cells containing a complete paired TCR. k) Percentage of BAL T cells containing a complete paired TCR for each participant across donor groups. l) Percentage of BAL T cells classified as TCR stimulated among cells containing a complete paired TCR. Each point represents an individual participant; boxes indicate the median and interquartile range. P values were calculated using the Kruskal–Wallis test, with pairwise comparisons performed using Dunn’s test with Benjamini–Hochberg (BH) correction. m) UMAP of BAL T cells colored according to TCR clonal expansion status. Clonotypes were classified as single (1 cell), small (2–5 cells), medium (6–20 cells), large (21–100 cells), or hyperexpanded (>100 cells).

Cluster 8 preferentially expressed effector cytokines (*IFNG, TNF, IL2*), inflammatory chemokines (*CCL3, CCL4*), cytotoxic molecules (*GZMB*), and activation-associated genes (*NFKBIA, TNFAIP3, CXCL9, CXCL10*), consistent with a highly activated antigen-responsive T-cell population. To distinguish TCR-dependent activation from bystander responses, TCR-Antigenic-Recognition-sequencing (TAR-seq) [20] was applied following Mtb stimulation (Figure 5f,g). TAR-seq-positive cells localized almost exclusively to cluster 8, whereas TAR-seq-negative cells were distributed across the remaining clusters, identifying cluster 8 as the principal Mtb-responsive T-cell population within BAL. TAR-seq additionally enabled high-confidence identification of putative Mtb-specific clonotypes for downstream analyses.

Mtb-responsive T cells represented only a small proportion of pulmonary T cells across all participant groups (Figure 5h,i). Although their frequency increased from community controls to PET^POS^ and active TB, this trend was not significant (p=0.554). PET^POS^ and active TB contained the highest proportions of Mtb-responsive cells, whereas community controls and converters contained relatively few. These findings indicate that most pulmonary T cells are not directly responding to Mtb antigen *ex-vivo*, despite increasing antigen-responsive populations in PET^POS^ and active TB. Mapping clonotype expansion onto the integrated UMAP demonstrated that singleton clonotypes were broadly distributed, whereas medium, large, and hyperexpanded clonotypes preferentially localized to cluster 8 (Figure 5k). Thus, the Mtb-responsive cluster is characterized by both TCR-dependent activation and substantial clonal expansion, consistent with antigen-driven proliferation.

### Conventional activation markers, PD-1, and interferon responses incompletely identify Mtb-responsive T cells

CD69 and CD154 are commonly used activation-induced markers (AIMs) [35–37], but failed to selectively identify the TAR-seq-defined Mtb-responsive population (Supplemental Figure 7a,b). Instead, CD69 and CD154 cells were distributed across multiple transcriptional clusters, demonstrating the improved specificity of TAR-seq for identifying TCR-dependent activation. Similarly, PD-1 expression was detected on a subset of pulmonary T cells but was broadly distributed across stimulated and unstimulated cells without enrichment following Mtb stimulation (Supplemental Figure 7c,d), indicating that PD-1 alone does not distinguish Mtb-responsive T cells. In addition to the TAR-seq-defined population, Mtb stimulation induced an interferon-stimulated T-cell population lacking evidence of TCR activation, consistent with cytokine-mediated bystander activation (Supplemental Figure 7e,f) [38,39].

### PET-CT identifies individuals at highest risk of progression to active TB and provides a natural model of protective immunity

All HHCs and CCs were followed prospectively for three years without preventive therapy (Figure 6a). Four participants developed microbiologically confirmed TB, all of whom had baseline PET^POS^ scans, whereas no PET^NEG^ or CC progressed. These findings demonstrate that PET-CT identifies individuals at markedly increased short-term risk of active TB. Among the 13 PET^POS^ participants, four (30.8%) progressed to microbiologically confirmed TB, whereas nine (69.2%) remained disease-free over three years. Because both groups had baseline PET-CT abnormalities despite negative microbiology, their divergent outcomes likely reflect differences in host immunity rather than disease stage. BAL T cells from progressors and non-progressors were therefore integrated and reclustered to identify transcriptional programs associated with progression or durable control (Figure 6b,c).

**Figure 6.**
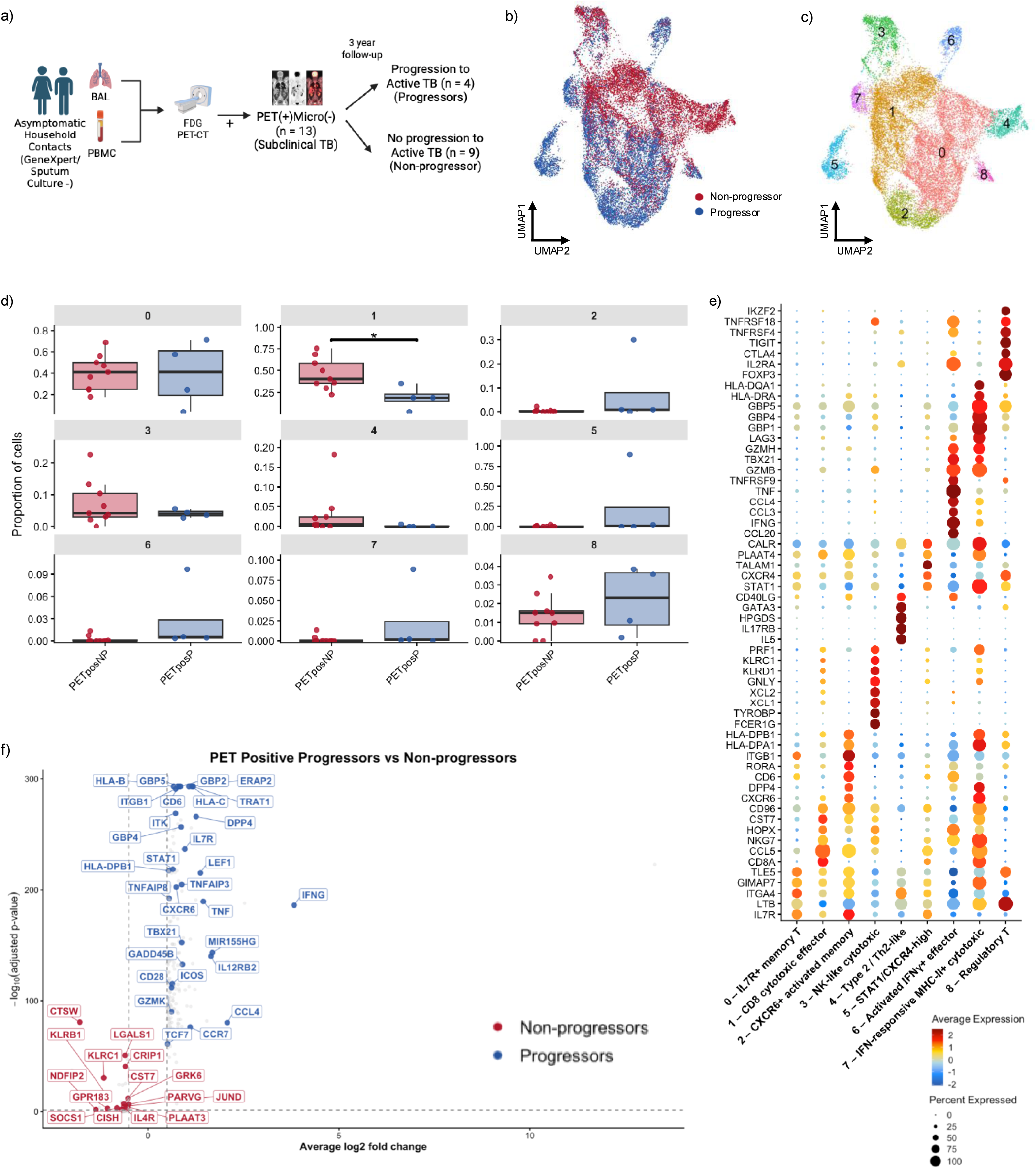
Pulmonary cytotoxic CD8 T-cell responses are associated with durable control of asymptomatic pulmonary tuberculosis. a) Study design. HHC and CC partipants were followed for 3 years and classified as “progresors” or “non-progressors based on development of subsequrnt microbiologciall confirmed TB. Progression only occurred in those from initial PET^POS^ group. b) UMAP of integrated BAL T cells from PET-positive progressors and non-progressors colored by clinical outcome. c) UMAP of BAL T cells colored by unbiased Leiden clusters. d) Proportion of cells within each cluster in PET-positive progressors and non-progressors. Each point represents an individual participant; boxes indicate the median and interquartile range. P values were calculated using the Kruskal–Wallis test, with pairwise comparisons performed using Dunn’s test with Benjamini–Hochberg (BH) correction. e) Dot plot showing expression of key marker genes used to annotate BAL T-cell clusters. Dot size represents the percentage of cells expressing each gene and color indicates scaled average expression. f) Volcano plot showing differentially expressed genes between BAL T cells from PET-positive progressors and non-progressors. Differential expression was performed using the Wilcoxon rank-sum test with Benjamini–Hochberg (BH) correction. Selected genes with an adjusted *P* < 0.05 and absolute log fold change > 0.5 are labeled.

### Individuals who durably control asymptomatic pulmonary tuberculosis are enriched for a cytotoxic CD8 T-cell population

Comparison of cluster abundance identified selective enrichment of cluster 1 in non-progressors (Figure 6d). This cluster displayed a canonical cytotoxic CD8 T-cell program characterized by *CD8A, CCL5, NKG7, KLRK1, KLRD1, HOPX,* and *ITGA1*, consistent with differentiated tissue-resident effector cells (Figure 6e; Supplemental Table 11). Their enrichment in non-progressors implicates pulmonary cytotoxic CD8 T cells as candidate mediators of protection. Differential expression analysis further demonstrated that non-progressors preferentially expressed cytotoxic genes (*CTSW, CST7, KLRC1, KLRB1, LGALS1*) together with activation-associated regulators (*CISH, JUND, JUNB*), whereas progressors upregulated inflammatory genes including *IFNG, TNF,* and *IFI6* (Figure 6f; Supplemental Table 12), suggesting that durable control is associated with cytotoxic rather than inflammatory T-cell programs.

### Isolation of pulmonary Mtb-reactive T-cell clones identifies candidate protective TCRs

To define the antigen specificity of pulmonary T cells responding to Mtb *in-vivo*, we generated 20 BAL-derived CD4 T-cell clones by limiting dilution (Figure 7a; Table 2). Paired single-cell TCR sequencing identified complete TCRα/β sequences for every clone, enabling direct linkage of TCR sequence with antigen specificity. Clones were screened against Mtb lysate, cell wall, culture filtrate proteins, MTB300, and the IMPAc-TB peptide library (Figure 7b–d) [40].

**Figure 7.**
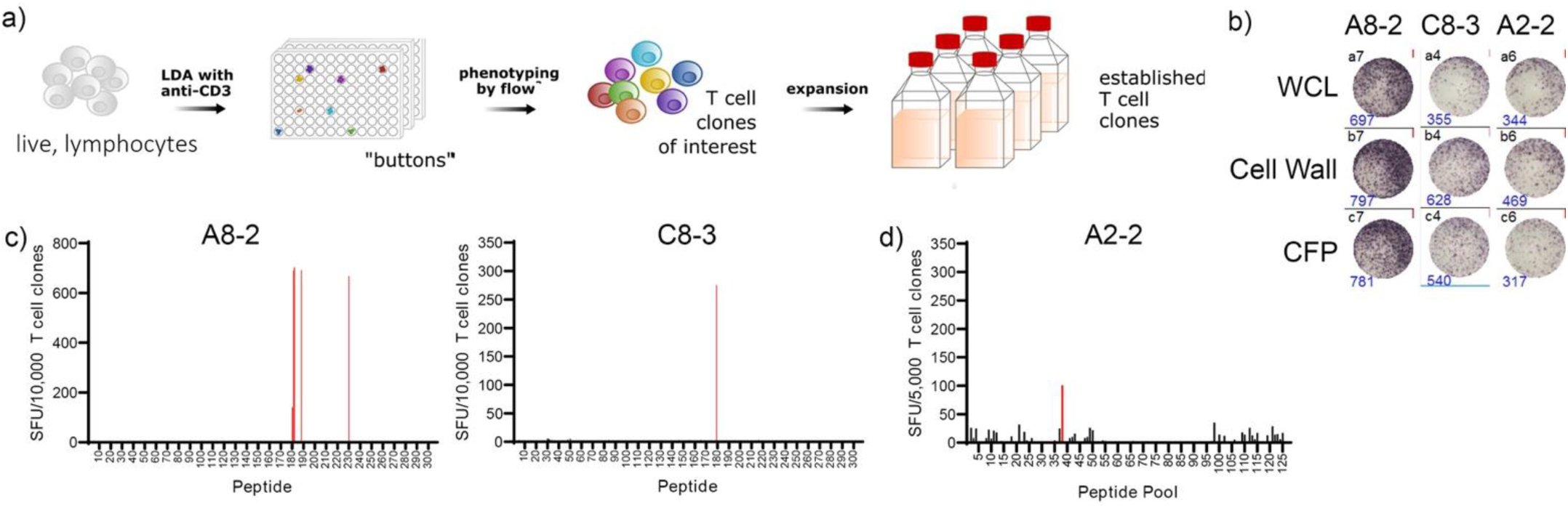
Limited Dilution Cloning from Lung T cells of an Active TB Participant. a) Live lymphocytes were sorted from BAL of an active TB participant. Limiting dilution cloning was performed with anti-CD3. Wells with growth after 2 weeks (buttons) were characterized by flow cytometry and then T cell clones of interest were expanded and characterized. b) T cell clones (10,000 cells/well) were incubated overnight with autologous LCL (10,000 cells/well) and either whole cell lysate (WCL), Cell Wall, and Culture Filtrate Protein (CFP) from Mtb in an IFN-γ ELISPOT. c) T cell clones (10,000 cells/well) were incubated overnight with autologous LCL (10,000 cells/well) and each individual peptide from the MTB300 peptide pool in an IFN-γ ELISPOT. d) T cell clones (5,000 cells/well) were incubated overnight with autologous LCL (5,000 cells/well) and the IMPAc-TB peptide library (127 wells) in an IFN-γ ELISPOT.

**Table 2:**
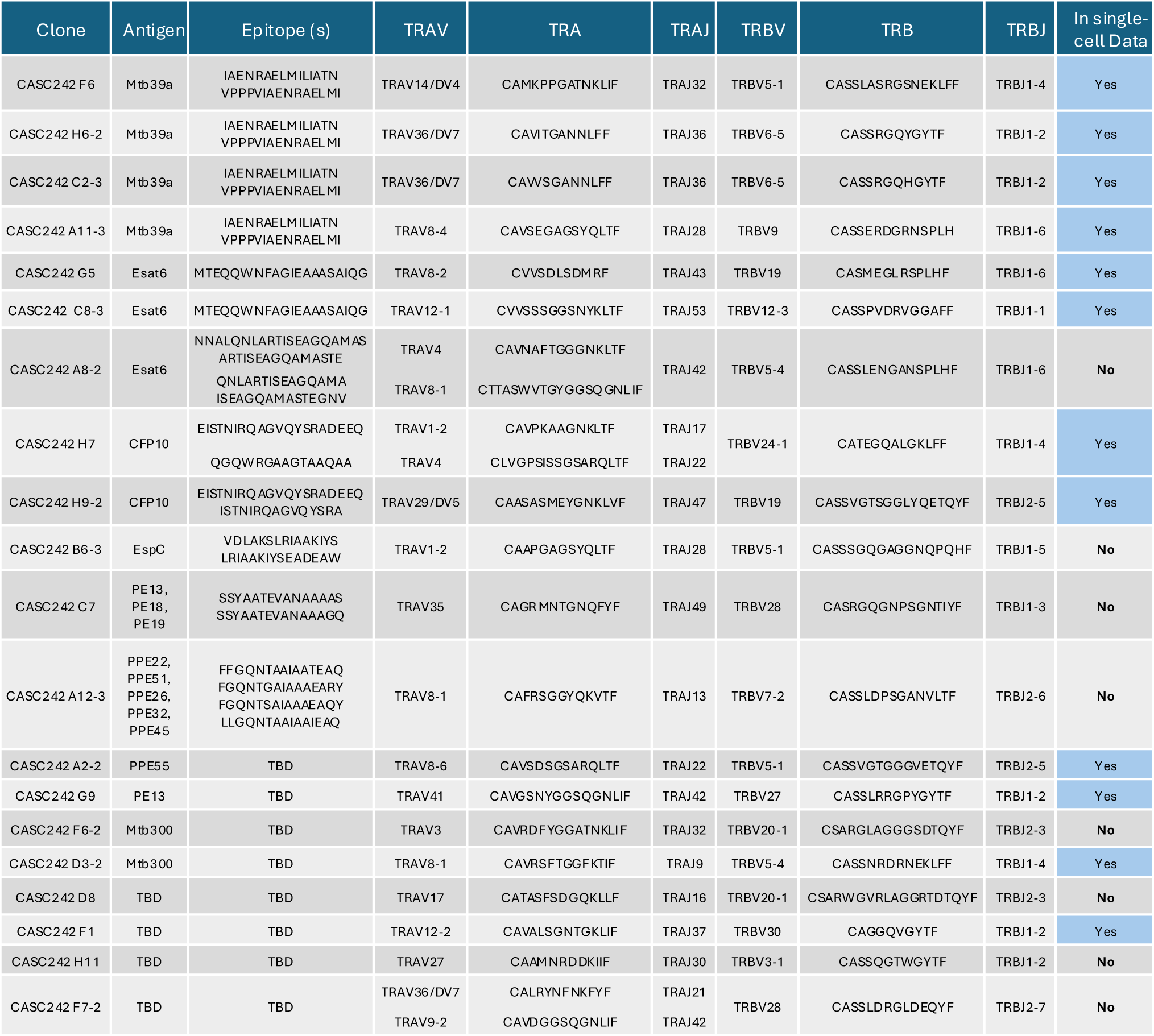
Paired TCR sequences and antigen specificities of BAL-derived T-cell clones from an individual with active TB. Twenty BAL-derived T-cell clones generated by limiting dilution from an active TB participant were functionally screened against Mtb antigens. The table summarizes clone identity, cognate antigen specificity, paired TCRα and TCRβ V/J gene usage and CDR3 sequences, and whether each clonotype was identified in the paired sc-TCR-seq dataset.

Recognized antigens included ESAT-6, CFP-10, Mtb39a, EspC, PPE55, PE13, PE13/18/19 family peptides, and other PE/PPE antigens, whereas four clones recognized Mtb but not any screened antigen (Table 2). These findings demonstrate broad antigenic diversity within the pulmonary T-cell response and establish a resource linking human pulmonary TCR sequences with defined Mtb antigen specificity.

### Antigen-specific pulmonary T-cell clones are recovered within the ex-vivo lung repertoire and undergo antigen-driven transcriptional remodeling

Exact paired TCRα/β matches were identified for 12 of 20 (60%) cloned TCRs within the independent single-cell dataset (Figure 8a), confirming that many functionally validated clonotypes are naturally represented *in-vivo*. Before stimulation, antigen-specific clonotypes occupied multiple transcriptional states (Figure 8b,c). Following stimulation, several—including ESAT-6-, PE13-, and Mtb39a-reactive clones—transitioned into the Mtb-responsive activation cluster, demonstrating rapid antigen-driven transcriptional remodeling.

**Figure 8.**
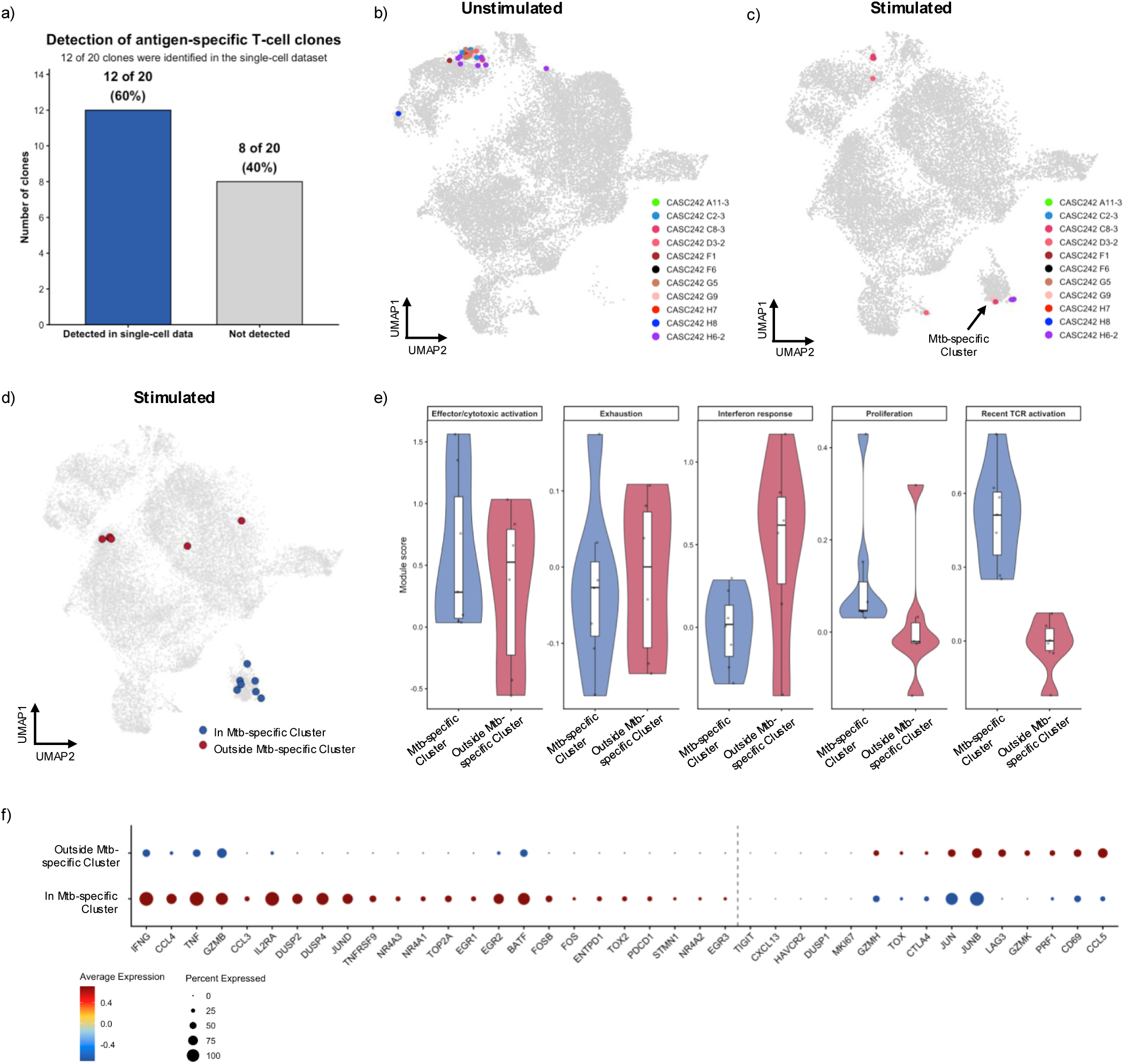
Antigen-specific pulmonary T-cell clonotypes undergo dynamic transcriptional remodeling following Mtb stimulation. a) Detection of functionally validated Mtb-specific T-cell clones within the paired single-cell TCR sequencing dataset. Bars indicate the number and percentage of cloned TCRs identified in the ex vivo BAL single-cell dataset. b) UMAP showing the distribution of functionally validated Mtb-specific T-cell clonotypes in unstimulated and Mtb-stimulated BAL T cells. Cells sharing identical paired TCRα/TCRβ sequences with cloned T cells are highlighted by clonotype. c) UMAP showing the distribution of the same antigen-specific clonotypes according to TAR-seq classification (TCR stimulated or not TCR stimulated). d) UMAP of stimulated BAL T cells showing the distribution of cells sharing identical paired TCRα/TCRβ sequences with clonotypes present in the Mtb-responsive cluster. Cells are classified according to whether they remain within or transition out of the Mtb-responsive cluster following stimulation. e) Comparison of transcriptional module scores between stimulated cells that remained within the Mtb-responsive cluster and clonally identical cells that transitioned to other transcriptional states. Violin plots show exhaustion, effector activation, proliferation, interferon response, and recent TCR activation module scores. P values were calculated using the Wilcoxon rank-sum test with Benjamini–Hochberg (BH) correction. f) Dot plot showing differential expression of activation-, effector-, proliferation-, and exhaustion-associated genes between stimulated cells that remained within the Mtb-responsive cluster and clonally identical cells that transitioned to other transcriptional states. Genes are ordered from those preferentially expressed in cluster-retained cells to those preferentially expressed in cells that transitioned to other transcriptional states. Dot size represents the percentage of cells expressing each gene and color indicates scaled average expression. Differential expression was performed using the Wilcoxon rank-sum test with Benjamini–Hochberg (BH) correction.

To determine whether entry into the Mtb-responsive transcriptional cluster was a stable feature of antigen-specific T cells, all cells sharing identical paired TCRα/TCRβ sequences with clonotypes present in the Mtb-responsive cluster were mapped across the integrated single-cell dataset (Figure 8d). Although most localized within the Mtb-responsive cluster, a subset occupied alternative transcriptional states. Compared with these cells, those within the Mtb-responsive cluster exhibited higher effector activation, cytotoxicity, proliferation, and TCR activation scores, whereas cells outside the cluster showed increased interferon-response and exhaustion-associated programs together with reduced proliferation and TCR activation (Figure 8e). Differential expression analysis supported these findings: cells within the Mtb-responsive cluster preferentially expressed activation and effector genes (*TNFRSF4, IL2RA, CCL20, CSF2, IFNG, TNF,* and *GZMB*), whereas cells outside the cluster expressed interferon-associated genes (*IFITM1, STAT1, IRF1, ISG15,* and *GBP5*) (Figure 8f). Together, these findings demonstrate that identical Mtb-specific clonotypes can occupy distinct functional states, with only a subset adopting a highly activated effector phenotype following antigen stimulation.

## Discussion

In this study, we provide, a comprehensive single-cell characterization of human pulmonary T-cell responses across the spectrum of Mtb exposure, asymptomatic infection, and active TB, while simultaneously defining the antigen specificity of lung-derived T cells. By integrating paired BAL and PBMC single-cell transcriptomics, TCR-sequencing, CITE-seq, longitudinal clinical follow-up, TAR-seq, and functional T-cell cloning, we demonstrate that pulmonary T-cell immunity is highly compartmentalized and reflects recent Mtb exposure. By following participants for 3 years, only those with initial PET^POS^ progressed to active TB. Those PET^POS^ who did non-progress, and were therefore protected, had an enrichment of cytotoxic CD8 T cells. Finally, by functionally defining pulmonary *Mtb*-specific TCRs and mapping these clonotypes within the single-cell atlas, we found that most BAL T cells were not *Mtb*-specific, while directly linking those that were to clonal identity, transcriptional state, and clinical outcome. Collectively, these findings provide new insights into the pulmonary immune mechanisms that govern successful containment of Mtb and identify candidate cellular and antigenic targets for next-generation TB vaccines.

One of the central findings of this study is that pulmonary T-cell immunity cannot be inferred from analyses of peripheral blood. Across every stage of Mtb exposure and disease, we observed marked compartmentalization of both transcriptional programs and TCR repertoires between the lung and circulation. Even within the same individual, relatively few clonotypes were shared between BAL and matched blood compartments, and those that were shared often demonstrated markedly different degrees of clonal expansion. These observations suggest that local antigen exposure within the lung drives selective recruitment, retention, and expansion of distinct T-cell populations, emphasizing that immune mechanisms operating at the site of infection are fundamentally different from those measured in blood. These observations are particularly important in the context of TB vaccine development where most candidate vaccines are evaluated using peripheral blood immunological readouts, yet these measurements have consistently failed to identify robust correlates of protection in clinical trials [41–43]. Our findings further support the need to define correlates of protective immunity through direct interrogation of pulmonary, rather than circulating, immune responses [44]. Such insights will be critical for the rational design of next-generation TB vaccines [44,45].

Our findings also indicate that the BAL T-cell compartment is distinct at different stages across the spectrum of TB infection and disease, without evidence of a clear progressive gradient. Instead, each clinical group displayed a distinct combination of T-cell states, suggesting that the pulmonary immune response evolves dynamically as exposure progresses toward infection and disease. We had anticipated that PET^NEG^ participants would resemble CC; instead, CC were distinguished by increase naïve and stem-like gene expression. This downregulation in PET^NEG^ suggests that an early immunological consequences of Mtb exposure occur locally within the lung, preceeding both radiographic inflammation and systemic sensitization. HHCs more broadly also exhibited reduced CCR7 expression, consistent with acquisition of a tissue-resident program, as CCR7 downregulation promotes lung retention and tissue-resident memory T-cell formation [29,46]. Accordingly, our findings suggest that pulmonary T cells in CC retain greater recirculatory potential, whereas *Mtb* exposure—even before PET-CT abnormalities or IGRA conversion—initiates tissue retention within the lung. The enrichment of activated pulmonary T cells in active TB is biologically plausible and consistent with sustained antigen exposure. Similar activated, cytotoxic, dysfunctional, and inflammatory T-cell states have been reported in active TB and in individuals progressing toward disease [19,47]. Rather than indicating protective immunity, these states likely reflect persistent antigenic stimulation and failure to achieve bacterial control. Accordingly, highly activated pulmonary T cells may represent a vigorous but ultimately ineffective immune response associated with ongoing inflammation and tissue injury [48].

A major barrier to defining protective immunity in human TB is that conventional tests cannot distinguish *Mtb* exposure, persistent infection, or incipient progression [49,50]. Consequently, protection has largely been inferred from peripheral blood rather than studied directly in early pulmonary disease with known clinical outcomes. PET-CT addresses this gap by identifying metabolically active pulmonary abnormalities before symptom onset or microbiological confirmation [51]. Longitudinal studies have established PET-CT abnormalities as markers of increased risk for active TB, consistent with biologically active preclinical disease [51–53].

Similarly, all incident TB cases in our cohort arose from participants with baseline PET-CT abnormalities, whereas no PET^NEG^ or CC participants progressed within 3 years. Although unsuitable for population screening, PET-CT provides a powerful platform to investigate protective immunity before clinical disease. In particular, PET^POS^ non-progressors exhibited pulmonary abnormalities compatible with asymptomatic TB yet remained disease-free for three years without treatment, suggesting effective endogenous immune control. Comparison with PET^POS^ progressors therefore enables protective immune mechanisms to be studied in individuals with similar baseline pulmonary abnormalities but divergent clinical outcomes.

Strikingly, non-progressors and progressors could be readily distinguished in our study. The enrichment of cytotoxic CD8 T-cell programs among non-progressors is particularly notable in this context. Conventional CD8 T cells recognize Mtb-infected cells through MHC class I, produce IFN-γ and TNF, and mediate perforin- and granzyme-dependent cytotoxicity [4,54].

They have been implicated in protection against TB in both experimental models and human disease [55,56]. More recently, studies in non-human primates have demonstrated that depletion of CD8α-expressing cells worsens TB disease [57] and abrogates protection induced by intravenous BCG vaccination, highlighting a broader role for CD8α-expressing cytotoxic lymphocytes that includes both conventional and unconventional T-cell populations [58,59]. Our findings suggest that a similar biology may operate in humans, with enrichment of a pulmonary *CD8A*-associated cytotoxic transcriptional program associated with durable control of asymptomatic pulmonary TB. Although our study does not distinguish the relative contributions of conventional and unconventional CD8α-expressing cells, it raises the possibility that both natural and vaccine-induced protection converge on shared cytotoxic immune mechanisms operating directly within the lung. These findings should nevertheless be interpreted cautiously. Only four participants progressed to active TB, and the identified protective signature will require validation in larger prospective PET-CT cohorts with longitudinal pulmonary sampling and functional characterization of the implicated cells and clonotypes. Nevertheless, the combination of baseline PET-CT, direct lung sampling, and prospective clinical follow-up provides a unique framework for defining pulmonary immune mechanisms associated with durable control of Mtb infection.

TAR-seq more specifically identified *Mtb*-responsive pulmonary T cells than conventional activation-induced marker (AIM) assays. Although CD69 and CD154 are widely used to detect antigen-responsive T cells [35–37], these markers reflect generalized activation rather than antigen-dependent TCR signaling. Accordingly, CD69 and CD154 cells spanned multiple transcriptional states and incompletely captured the discrete *Mtb*-reactive population defined by TAR-seq [20]. By capturing transcriptional programs induced by recent TCR engagement, TAR-seq may therefore better resolve antigen-responsive T cells in highly activated tissues such as the lung.

A major challenge in TB immunology is identifying T-cell receptors that recognize physiologically infected cells, as peptide-reactive T cells do not necessarily recognize naturally infected macrophages [60,61]. By coupling TAR-seq with paired TCR sequencing and functional cloning, we directly linked pulmonary TCR sequences to Mtb antigen specificity, transcriptional phenotype, and clonal architecture. Most cloned TCRs were independently identified in the *ex-vivo* single-cell dataset, providing orthogonal validation of this approach. Finally, identical antigen-specific clonotypes occupied multiple transcriptional states within the lung. Although many transitioned into the TAR-seq-defined activation state following stimulation, clonally identical cells also exhibited quiescent, interferon-conditioned, and exhaustion-associated programs with reduced proliferative capacity. These findings indicate that antigen specificity alone does not determine T-cell function and suggest that regulation of T-cell state may be as important as antigen recognition in protective immunity.

These findings also have important implications for TB vaccine development. Rather than simply increasing the frequency of antigen-specific T cells, effective vaccines may need to establish pulmonary tissue-resident T cells capable of rapidly transitioning into highly functional cytotoxic effector states following pathogen encounter [41]. By integrating TAR-seq with paired TCR sequencing and functional cloning, our approach provides a platform for identifying these protective clonotypes and the naturally processed antigens they recognize, informing rational antigen selection for next-generation TB vaccines. By integrating longitudinal clinical phenotyping with direct interrogation of the human lung, our study links transcriptional state, antigen specificity, clonal identity, and clinical outcome in tuberculosis. These findings redefine pulmonary T-cell immunity across the spectrum of Mtb exposure and disease, identify candidate immune mechanisms associated with durable control of asymptomatic infection, and provide a framework for discovering the protective T-cell responses and antigens that will inform the rational design of next-generation TB vaccines.

## Methods

### Study Participants

This study was conducted according to the principles expressed in the Declaration of Helsinki. Study participants, protocols, and consent forms were approved by the Institutional Review Board of the University of Stellenbosch (IRB00000186; N19/10/150).

Written informed consent was obtained from all participants before enrollment. All ethical regulations relevant to human research participants were followed. All participants were from the Cape Town region, South Africa. Fully study protocol is previously published [21]. Briefly, those with active microbiologically confirmed (GeneXpert or culture positive) pulmonary TB were enrolled from the Cape Town area. Then their HHCs with a minimum 3 months of exposure, but who remained asymptomatic and whose sputum remained negative by GeneXpert and culture were enrolled. HHCs had IGRA placed at time of enrollment and 3 months later, as well as undergoing an 18F-Fluorodeocyglucose PET-CT scan to further delineate those with IGRA positive at month 0 and 3 with negative PET-CT scan (PET^NEG^), those with IGRA negative at month 0 and positive at month 3 but with negative PET-CT scan (Converter), and those with positive PET-CT scan (PET^POS^). Additionally community control participants from the same communities but without known recent exposures were enrolled, who were all IGRA positive at baseline and also all had negative PET-CT scan. All CC and HHC participants were followed for 3 years for the development of microbiologically confirmed TB.

### Collection of BAL and PBMC Samples

Human monocyte-derived dendritic cells (DCs) were isolated from Peripheral blood mononuclear cells (PBMC) by plate adherence. For this, PBMCs were incubated in RPMI supplemented with 2% heat-inactivated human serum, 3.5 mM L-Glutamine (Gibco), and 44 μg/ml gentamycin (Gibco) at 37°C for 1h, before gentle tapping and removal of non-adherent cells. Adherent cells were then cultured in RPMI (Gibco) supplemented with 10% heat-inactivated human serum (RPMI 10%HuS) and 30 ng/ml GM-CSF (Immunex) and 10 ng/ml IL-4 (R&D Systems) for 5 days [62].

*M. tuberculosis (Mtb) auxotroph Stimulation: M. tuberculosis* (Mtb) auxotroph mc^2^6206 (H37Rv ΔpanCD ΔleuCD) [22] was as a kind gift from Bill Jacobs (MTA-IN19-168) and is modified such that it is only able to grow in the presence of externally supplemented pantothenate and leucine and thus can be handled in a biosafety level 2 setting. The Mtb auxotroph was grown to OD_600_ 0.5 in 7H9 broth supplemented with Leucine (50mg/ml), Pantothenate (24mg/ml), and 5%

Tween 80. 1e5 DCs were either uninfected or infected with Mtb auxotroph at MOI 9 and incubated overnight. PBMC and non-adherent BAL were thawed in the presence of DNase and initially resuspended in RPMI10%HuS. T cells were isolated from the PBMCs and BAL in one of two ways. In the first method, CD3+ T cells were isolated using negative selection with the Pan T cell Isolation Kit (Miltenyi Biotec, 130-096-535) per the manufacturer’s instructions. BAL underwent an additional lymphocyte enrichment via CD11b depletion using CD11b microbeads (Miltenyi Biotec, 130-049-601) per the manufacturer’s instructions. In the second method, cells were resuspended in RPMI10%HuS at a concentration of 1e7/ml. Propidium iodide (PI; Miltenyi, 130-093-233) was added at 0.25μl/2e6 cells. Samples were then sorted using a BD Influx™ Cell Sorter for live (PI negative) lymphocytes (Supplemental Figure 1c). Mtb-infected and uninfected DCs (1e4) were then incubated with T cells (1e5) overnight. Cells were incubated in Human TruStain FcX (Biolegend^TM^) for 10 minutes at 4°C and then stained with TotalSeq-C hashtag antibodies and CITE-seq antibodies (Biolegend^TM^) listed in Supplemental Table 13 for 30 minutes at 4°C.

### Single-cell Sequencing of BAL and PBMC Samples

Cells were then loaded in either the10X Genomics^TM^ Chromium with Next GEM Single Cell 5’ v2 or the Chromium iX using the GEM-X Single Cell 5’ v3. Full protocol details of single-cell GEX, TCR and cell surface protein library preparation can be found from 10X genomics website (https://cdn.10xgenomics.com/image/upload/v1722286086/support-documents/CG000330_Chromium_Next_GEM_Single_Cell_5_v2_Cell_Surface_Protein_UserGuide_RevG.pdf or https://cdn.10xgenomics.com/image/upload/v1710231306/support-documents/CG000734_ChromiumGEM-X_SingleCell5_ReagentKitsv3_CellSurfaceProtein_UserGuide_RevA.pdf). Finished libraries were then sent to Novogene Corporation Inc. in Sacramento California for NovaSeq 6000 sequencing.

### Single-Cell RNA-seq Pre-processing

Raw sequence reads were processed using 10X Genomics Cell Ranger software (version 6.1.1). The resulting sequence data were aligned to the GRCh38 human genome. Cell demultiplexing used a combination of algorithms, including GMM-demux, demuxEM and BFF, implemented using the cellhashR package [63–65]. Droplets identified as doublets (i.e. the collision of distinct sample barcodes) were removed from downstream analyses. We additionally performed doublet detection using DoubletFinder, and removed doublets from downstream analysis [66]. Next, droplets were filtered based on UMI count (allowing 0-20,000/cell), and unique features (allowing 200-5000/cell). Additionally, we computed a per-cell saturation statistic for both RNA and ADT data, defined as: 1 – (#UMIs / #Counts). This statistic provides a per-cell measurement of the completeness with which unique molecules are sampled per cell and has the benefit of being adaptable across diverse cell types. Data were filtered to require RNA saturation > 0.35. Analyses utilized the Seurat R package, version 4.2 [67]. Using standardized methods implemented in the Seurat R package, counts and UMIs were normalized across cells, scaled per 10,000 bases, and converted to log scale using the ’NormalizeData’ function. These values were then converted to z-scores using the ’ScaleData’ command. Highly variable genes were selected using the ’FindVariableGenes’ function with a dispersion cutoff of 0.5. Principal components were calculated for these selected genes and projected onto all other genes using the ’RunPCA’ and ’ProjectPCA’ commands. Clusters of similar cells were identified using the Louvain method for community detection, and UMAP projections were calculated. CITE-seq data were CLR-normalized by lane, meaning raw count data are subset per lane, CLR normalization performed (as implemented in the Seurat R package, using margin = 1), using all QC-passing cells/lane. Normalization per-lane was performed to reduce batch effects.

### Gene Module Scores

The following published gene modules scores were used in the paper and their component genes are listed: interferon response modules score (*IFI6, IFI27, MX1, ISG15, STAT1, MX2, IFIT3*), cytotoxicity (*PRF1, GNLY, NKG7, GZMA, GZMB, GZMH, GZMK* and *GZMM*), glycolysis module score (*ALDOA, BPGM, ENO1, ENO2, GAPDH, HK1, HK2, HKDC1, PFKL, PGAM1, PGAM2, PGK1, PKLR, PKM, TPI1*) and naïve T cell module score (*CTSH, CA6, LEF1, RGS10, TMIGD2*) [26].

*TCR Sequence Analysis:* Raw sequence reads for gene expression and TCR enrichment were first processed using cellranger software, version 6.1.1 (10X Genomics). The raw clonotype calls produced by cellranger vdj were extracted from the comma-delimited outputs. Cells were demultiplexed and TCR calls were assigned to samples using custom software, made publicly available through the cellhashR package, with the GMM-Demux, demuxEM, and BFF algorithms [63–65]. The clonotype data generated by cellranger were filtered to drop any cells where the TCR calls lacked a CDR3 sequence, the clonotype was not marked as full-length, or the clonotype lacked a called V, J, or constant gene. Rows with chimeric V/J/C combinations (i.e. TRBV / TRAC) were filtered, with the exception that segments consisting of a TRDV/TRAC or TRAV/TRDC were permitted. These chimeric segments were classified according to the constant region chain. Data was analyzed in R studio using Seurat package [67].

All code is available on github (https://github.com/kaindylan/IMPAcTB---Lung-resident-T-Cells/tree/main).

### Limited Dilution Cloning and Flow Confirmation of Initial Phenotype

Non-adherent BAL samples were stained with a viability dye (Propidium Iodide, Miltenyi Biotec). Live cells were sorted by size using FSC/SSC gating to collect the lymphocyte population. The cells were plated in a 96-well plate in a limited dilution assay with 1.5×10^5^ irradiated PBMC (3000 cGray) and 3×10^4^ irradiated LCL (6000 cGray) along with αCD3 (30ng/ml), IL-2 (2ng/ml), IL-7 (0.5ng/ml), IL-12 (0.5ng/ml) and IL-15 (0.5ng/ml) in RPMI 1640 supplemented with 10% heat inactivated human serum in 96 well round bottom plates. On day 5, the αCD3 was washed out by removal of half of the volume from the wells and replaced with additional cytokine supplemented media.

Media was changed for all wells every 2-3 days and replaced with cytokine supplemented media. T cell “buttons” were evaluated for growth on Day 20 and selected buttons stained for fluorescence bar coding using Cell Trace Far Red (Thermofisher) and Cell Trace Violet (Thermofisher) as well as a phenotyping panel that included the MR1/5-OP-RU tetramer (NIH Tetramer Core), CD3 PE-Cy7 (clone SK7; Biolegend), CD4 BUV737 (clone SK3; Waters Biosciences), CD8 APC-Cy7 (clone SK1; Biolegend), TCRαβ BV605 (clone IP26; Biolegend), TCRγδ FITC (clone B1; Biolegend), TCR Vα24-Jα18 BV711 (clone 6B11; Biolegend).

### Statistics

Statistical analyses were performed in R (v4.4.2) unless otherwise specified. Single-cell differential gene expression analyses were performed using the Wilcoxon rank-sum test implemented in Seurat, with Benjamini–Hochberg (BH) correction for multiple testing. Genes with an adjusted *P* value <0.05 were considered significantly differentially expressed. Additional log_2_fold-change thresholds are indicated in the corresponding figure legends where applied. Pseudobulk differential expression analyses were performed using DESeq2 with BH correction for multiple testing, with an adjusted *P* value <0.05 considered statistically significant. Comparisons of cell proportions, module scores, CITE-seq protein expression, TCR diversity, and clonotype frequencies between independent groups were performed using the Kruskal–Wallis test followed by Dunn’s post hoc test with BH correction for multiple comparisons. Paired comparisons between matched BAL and PBMC samples were performed using the paired Wilcoxon signed-rank test with BH correction. Correlations between BAL and PBMC clonotype frequencies were assessed using Spearman’s rank correlation coefficient. Gene Ontology enrichment analyses were performed using clusterProfiler with hypergeometric testing and BH correction for multiple testing. Unless otherwise specified, two-sided *P* values <0.05 after multiple testing correction were considered statistically significant. Samples from household contacts and community controls were processed and analyzed in a blinded fashion with respect to tuberculosis infection and disease status.

### Data Availability

All R studio code used in the generation of this analysis is freely available and uploaded to github (https://github.com/kaindylan/IMPAcTB---Lung-resident-T-Cells/tree/main). All raw data was deposited into the Sequence Read Archive database. The authors declare that the data supporting the findings of this study are available within the paper and its supplementary information files. The source data underlying the graphs in the paper can be found in the Supplementary Data. Raw data is deposited on dbGAP.

## Author Contribution

DK, GW, NDP, DAL, BNB, DML contributed to the conception and/or design of the work. GW, NDP contributed to clinical activities. DML, DAL, BNB, GW, NDP received funding for the research. DK, GWM, KHR, MC, GMS, GW, NDP, DAL, BNB, DML substantially contributed to the acquisition, analysis, or interpretation of data and drafting of the manuscript. All authors substantially contributed to revising and critically reviewing the manuscript for important intellectual content. All authors approved the final version of this manuscript to be published and agree to be accountable for all aspects of the work.

## Acknowledgements

We would like to thank the BioMedical Research Institute (BMRI) Clinical Team and the Stellenbosch University Immunology Research Group laboratory team including Stephanus Malherbe, Andriëtte Hiemstra, Ilana van Rensburg, Marika Flinn, Ayanda Shabangu. We acknowledge the assistance of the Oregon Clinical & Translational Research Institute, which is supported by the National Center for Advancing Translational Sciences, National Institutes of Health, through Grant Award Number UL1TR002369. This project used the OHSU Flow Cytometry and Monoclonal Antibody Shared Resource Core Facility (RRID:SCR_009974). The research reported in this publication used computational infrastructure supported by the Office of Research Infrastructure Programs, Office of the Director, of the National Institutes of Health under Award Number S10OD034224.

## Supplemental

**Supplemental Figure 1: BAL Cell Differentials Are Unchanged Over the Spectrum of TB Infection/Disease**

a) Relative abundance of major BAL cell populations for each participant, including macrophages, lymphocytes, neutrophils, and eosinophils. Each bar represents an individual participant.

b) Proportion of macrophages, lymphocytes, neutrophils, and eosinophils in BAL across donor groups. Each point represents an individual participant; boxes indicate the median and interquartile range. Overall group comparisons were performed using the Kruskal–Wallis test, with Benjamini–Hochberg (BH)-adjusted *P* values.

c) Representative FACS sorting strategy.

**Supplemental Figure 2 – Distinct Transcriptional Profiles Distinguish BAL and PBMC T cells**

a) Absolute number of unstimulated T cells included in analysis from each participant’s BAL and PBMC. Boxes indicate the median and interquartile range; each point represents an individual participant.

b) Proportion of BAL and PBMC T cells within each unbiased Leiden cluster. Each point represents an individual participant; boxes indicate the median and interquartile range. P values were calculated using the Wilcoxon rank-sum test with Benjamini–Hochberg (BH) correction.

c) Volcano plot showing differentially expressed genes between BAL and PBMC T cells identified by pseudobulk differential expression analysis. Differential expression was performed using DESeq2 with Benjamini–Hochberg (BH)-adjusted *P* values. Genes with an adjusted *P* < 0.05 and absolute log fold change > 0.25 are labeled. Positive log fold changes indicate genes enriched in BAL T cells, whereas negative log fold changes indicate genes enriched in PBMC T cells.

**Supplemental Figure 3 – Pulmonary T-cell compartmentalization is preserved in CC participants.**

a) UMAP of T cells from community control participants colored by tissue of origin (BAL or PBMC).

b) UMAP of community control T cells colored by unbiased Leiden clusters.

c) Proportion of BAL and PBMC T cells within each Leiden cluster in community control participants. Each point represents an individual participant; boxes indicate the median and interquartile range. P values were calculated using the Wilcoxon rank-sum test with Benjamini–Hochberg (BH) correction.

d) Volcano plot showing differentially expressed genes between BAL and PBMC T cells from community control participants identified by pseudobulk differential expression analysis. Differential expression was performed using DESeq2 with Benjamini–Hochberg (BH)-adjusted *P* values. Genes with an adjusted *P* < 0.05 and absolute log fold change > 0.25 are labeled. Positive log fold changes indicate genes enriched in BAL T cells, whereas negative log fold changes indicate genes enriched in PBMC T cells.

e) Comparison of transcriptional module scores between BAL and PBMC T cells from community control participants. Each point represents an individual participant; boxes indicate the median and interquartile range. P values were calculated using the Wilcoxon rank-sum test with BH correction.

f) Comparison of CITE-seq protein expression between BAL and PBMC T cells from community control participants. Each point represents an individual participant; boxes indicate the median and interquartile range. P values were calculated using the Wilcoxon rank-sum test with BH correction.

**Supplemental Figure 4 – Predominantly CD4 T-cell clusters exhibit distinct transcriptional changes across the spectrum of tuberculosis infection and disease.**

a) Classification of Leiden clusters according to CITE-seq-defined lineage as predominantly CD4, predominantly CD8, mixed, or double negative.

b-g) Volcano plots showing pseudobulk differential expression analysis of predominantly CD4 T-cell clusters between donor groups: b) Control versus PETNEG, c) Control versus Converter, d) Control versus PETPOS, e) Active TB versus Control, f) Active TB versus PETNEG, and g) Active TB versus Converter. Differential expression was performed using DESeq2 with Benjamini–Hochberg (BH)-adjusted *P* values. Genes with an adjusted *P* < 0.05 and absolute log fold change > 0.25 are labeled.

**Supplemental Figure 5 – Predominantly CD8 T-cell clusters exhibit distinct transcriptional changes across the spectrum of tuberculosis infection and disease.**

a–g) Volcano plots showing pseudobulk differential expression analysis of predominantly CD8 T-cell clusters between donor groups: a) Control versus PETNEG, b) Control versus Converter, c) Control versus PETPOS, d) PETPOS versus PETNEG, e) Converter versus PETNEG, f) Active TB versus Control, and g) Active TB versus PETPOS. Differential expression was performed using DESeq2 with Benjamini–Hochberg (BH)-adjusted *P* values. Genes with an adjusted *P* < 0.05 and absolute log fold change > 0.25 are labeled.

**Supplemental Figure 6 – TAR-seq more specifically identifies Mtb-responsive pulmonary T cells than conventional transcriptional signatures.**

a) UMAP of BAL T cells coloured by CD154 CITE-seq expression.

b) UMAP of BAL T cells coloured by CD69 CITE-seq expression.

c) UMAP of unstimulated BAL T cells coloured by PD-1 CITE-seq expression.

d) UMAP of stimulated BAL T cells coloured by PD-1 CITE-seq expression.

e) UMAP of unstimulated BAL T cells coloured by interferon response gene module (*IFI6, IFI27, MX1, ISG15, STAT1, MX2, IFIT3*).

f) UMAP of stimulated BAL T cells coloured by interferon response gene module (*IFI6, IFI27, MX1, ISG15, STAT1, MX2, IFIT3*).

