## Supplemental Table 13 for "Lung Resident T Cells Across the Spectrum of TB Infection and Disease"

| **TotalSeq Antibodies:** | **Manufacturer** | **Catalog number** |
| --- | --- | --- |
| CD4 | Biolegend | 300567 |
| CD8 | Biolegend | 344753 |
| CD45RA | Biolegend | 304163 |
| CD62L | Biolegend | 304851 |
| CD26 | Biolegend | 302722 |
| PD1/CD279 | Biolegend | 329963 |
| CD38 | Biolegend | 303543 |
| CD27 | Biolegend | 302853 |
| CD11b | Biolegend | 301359 |
| CD25 | Biolegend | 302649 |
| CD69 | Biolegend | 310951 |
| CD154 | Biolegend | 310849 |
| TotalSeq C0251 Anti-Human Hashtag #1 | Biolegend | 394661 |
| TotalSeq C0252 Anti-Human Hashtag #2 | Biolegend | 394663 |
| TotalSeq C0253 Anti-Human Hashtag #3 | Biolegend | 394665 |
| TotalSeq C0254 Anti-Human Hashtag #4 | Biolegend | 394667 |
| Human TruStain FcX (Fc Receptor Blocking Solution) | Biolegend | 422302 |

Supplemental Table 8.
