## Supplemental Figures for "Lung Resident T Cells Across the Spectrum of TB Infection and Disease"

a)

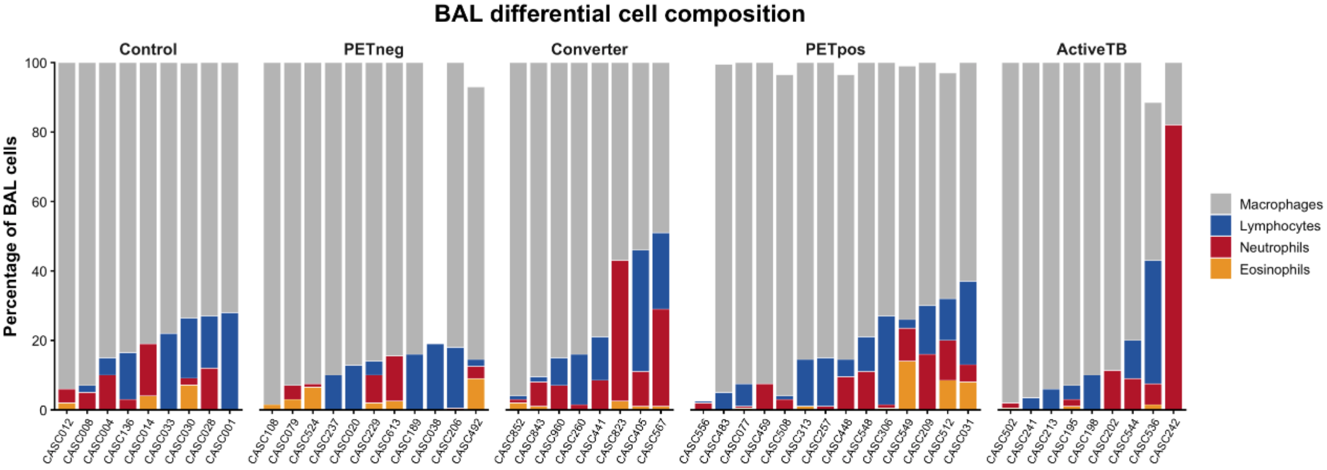
BAL differential cell composition
Control
PETneg
Converter
PETpos
ActiveTB
Percentage of BAL cells
0
20
40
60
80
100
Macrophages
Lymphocytes
Neutrophils
Eosinophils
CASC012 CASC008 CASC004 CASC136 CASC014 CASC033 CASC030 CASC028 CASC001
CASC108 CASC079 CASC524 CASC237 CASC020 CASC229 CASC613 CASC189 CASC038 CASC206 CASC492
CASC652 CASC843 CASC960 CASC260 CASC441 CASC823 CASC405 CASC567
CASC556 CASC483 CASC077 CASC459 CASC508 CASC313 CASC257 CASC448 CASC548 CASC306 CASC549 CASC209 CASC512 CASC031
CASC502 CASC241 CASC213 CASC195 CASC198 CASC202 CASC544 CASC536 CASC242

b)

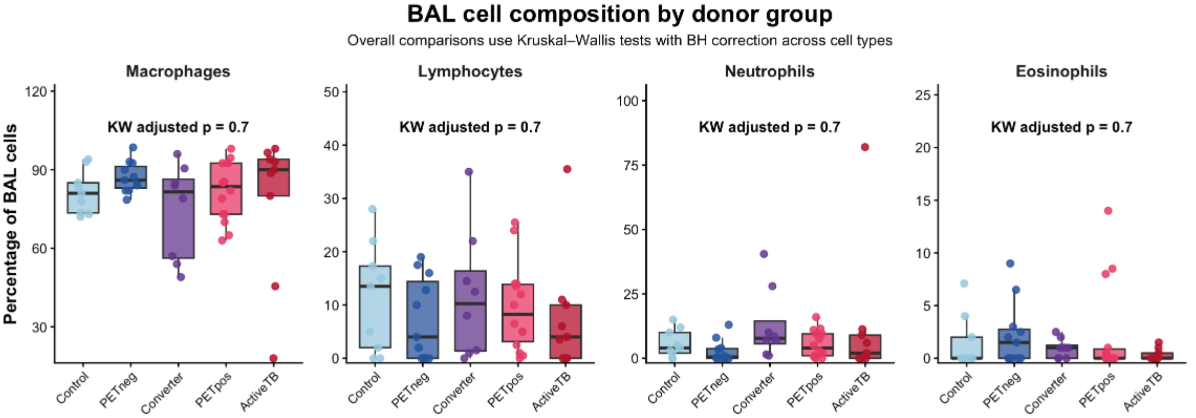
BAL cell composition by donor group
Overall comparisons use Kruskal–Wallis tests with BH correction across cell types
Macrophages
Lymphocytes
Neutrophils
Eosinophils
KW adjusted p = 0.7
KW adjusted p = 0.7
KW adjusted p = 0.7
KW adjusted p = 0.7
Percentage of BAL cells
Control
PETneg
Converter
PETpos
ActiveTB

c)

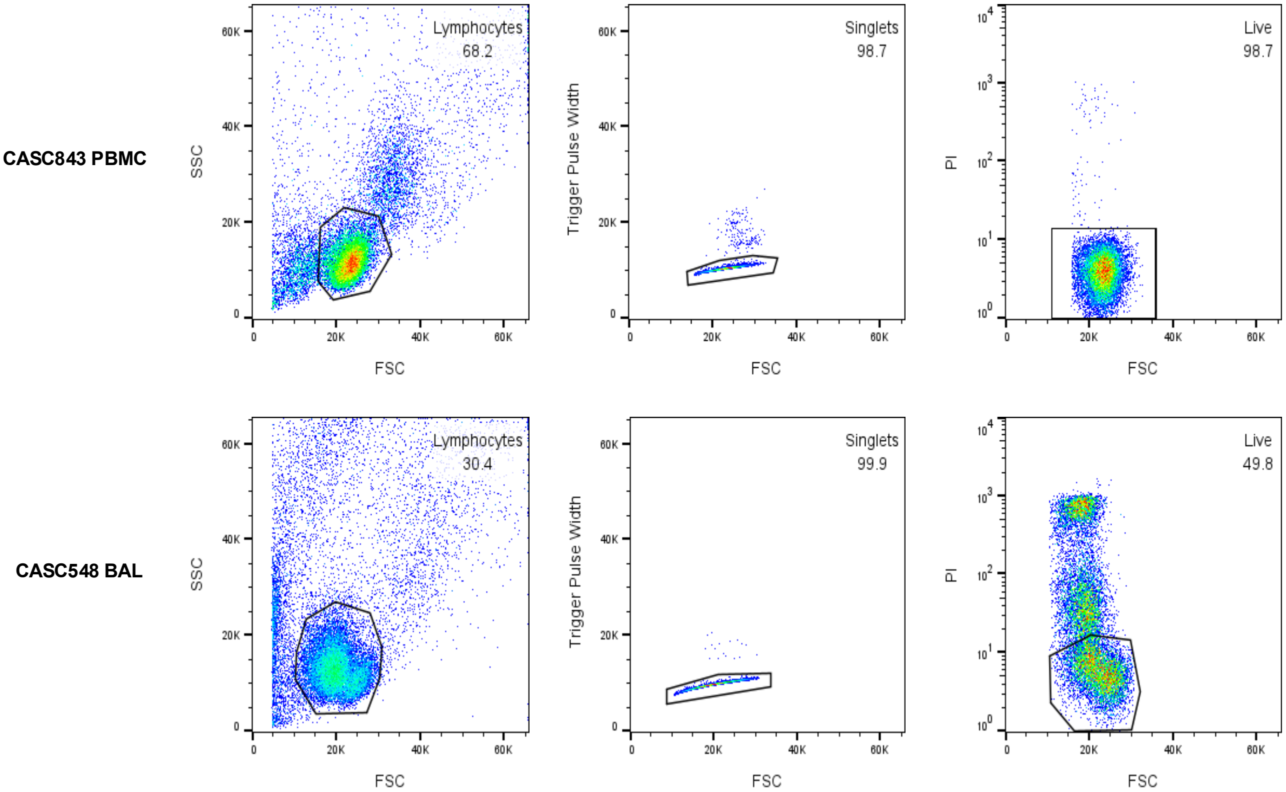
CASC843 PBMC
CASC548 BAL
Lymphocytes
68.2
Singlets
98.7
Live
98.7
Lymphocytes
30.4
Singlets
99.9
Live
49.8
SSC
Trigger Pulse Width
PI
FSC

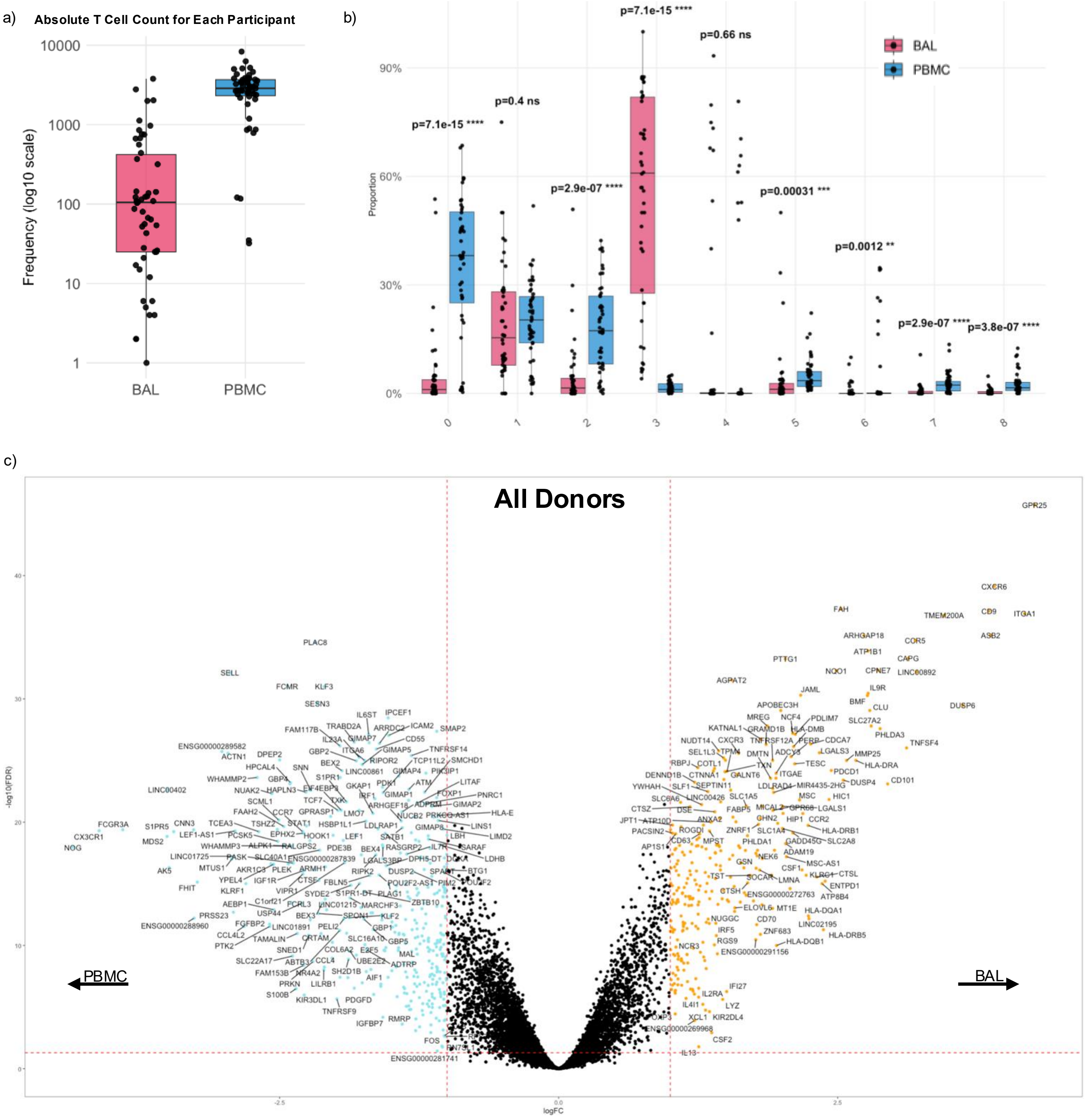

a)
Absolute T Cell Count for Each Participant
Frequency (log10 scale)
10000
1000
100
10
1
BAL
PBMC
b)
Proportion
90%
60%
30%
0%
BAL
PBMC
p=7.1e-15 ****
p=0.4 ns
p=2.9e-07 ****
p=7.1e-15 ****
p=0.66 ns
p=0.00031 ***
p=0.0012 **
p=2.9e-07 ****
p=3.8e-07 ****
0
1
2
3
4
5
6
7
8
c)
All Donors
-log10(FDR)
logFC
PBMC
BAL
GPR25
CXCR6
CD9
ITGA1
FAH
TMEM200A
ASB2
PLAC8
SELL
FCMR
KLF3
SESN3
CX3CR1
NOG
FCGR3A

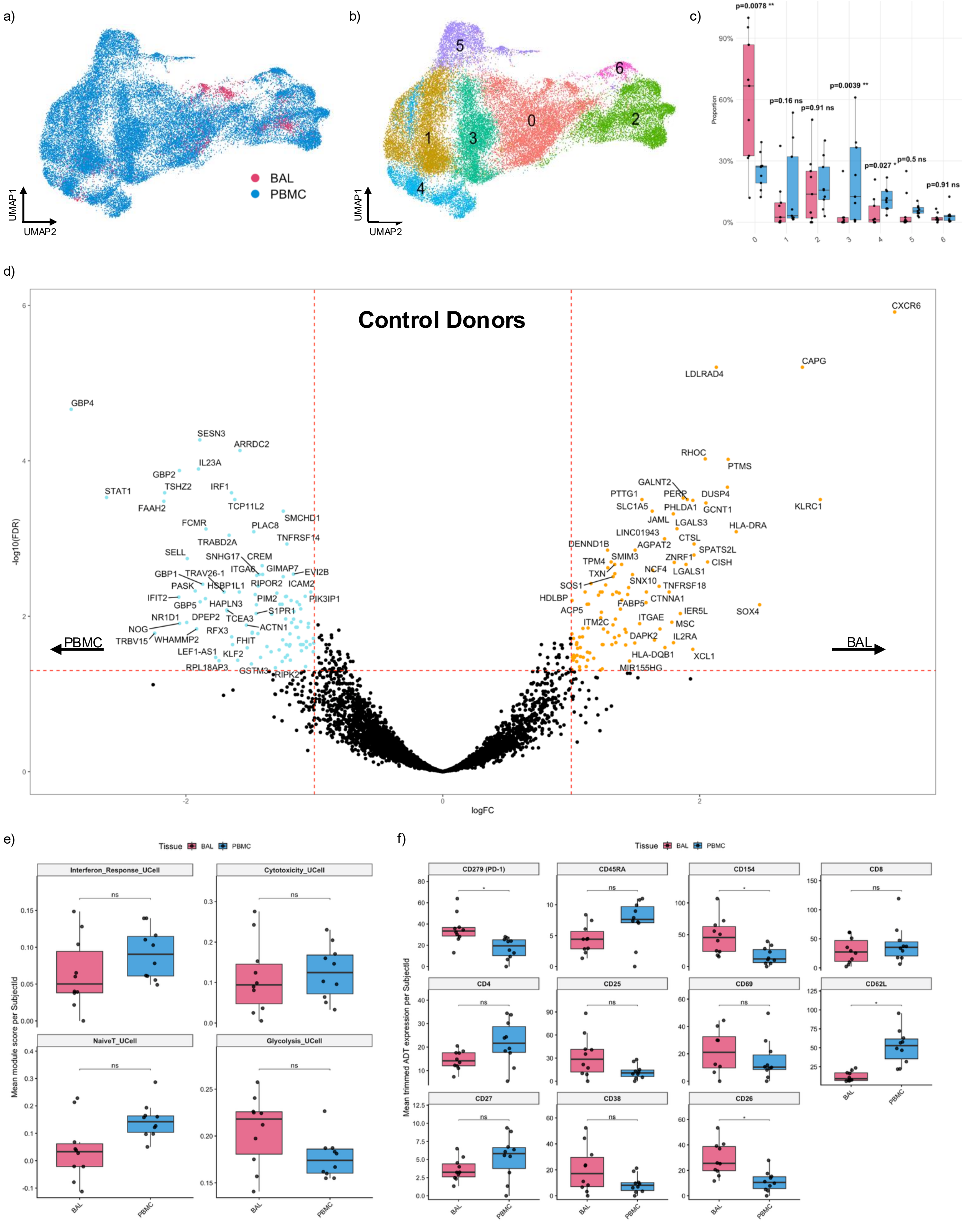

a)
BAL
PBMC
UMAP1
UMAP2
b)
5
6
1
3
0
2
4
UMAP1
UMAP2
c)
p=0.0078 **
p=0.16 ns
p=0.91 ns
p=0.0039 **
p=0.027 *
p=0.5 ns
p=0.91 ns
Proportion
90%
60%
30%
0%
0
1
2
3
4
5
6
d)
Control Donors
-log10(FDR)
logFC
PBMC
BAL
CXCR6
CAPG
LDLRAD4
GBP4
SESN3
ARRDC2
IL23A
GBP2
TSHZ2
IRF1
STAT1
FAAH2
TCP11L2
SMCHD1
FCMR
PLAC8
TNFRSF14
TRABD2A
SELL
SNHG17
CREM
ITGA6
GIMAP7
EVI2B
GBP1
TRAV26-1
PASK
HSBP1L1
RIPOR2
ICAM2
IFIT2
GBP5
HAPLN3
PIM2
PIK3IP1
S1PR1
NR1D1
DPEP2
TCEA3
NOG
RFX3
ACTN1
TRBV15
WHAMMP2
FHIT
LEF1-AS1
KLF2
RPL18AP3
GSTM3
RIPK2
RHOC
PTMS
GALNT2
PTTG1
PERP
DUSP4
SLC1A5
PHLDA1
GCNT1
KLRC1
JAML
LGALS3
HLA-DRA
LINC01943
CTSL
DENND1B
AGPAT2
SPATS2L
SMIM3
ZNRF1
TPM4
CISH
TXN
NCF4
LGALS1
SOS1
SNX10
TNFRSF18
HDLBP
CTNNA1
FABP5
ACP5
IER5L
SOX4
ITGAE
ITM2C
MSC
DAPK2
IL2RA
HLA-DQB1
XCL1
MIR155HG
e)
Tissue
BAL
PBMC
Interferon_Response_UCell
Cytotoxicity_UCell
NaiveT_UCell
Glycolysis_UCell
Mean module score per SubjectId
ns
f)
CD279 (PD-1)
CD45RA
CD154
CD8
CD4
CD25
CD69
CD62L
CD27
CD38
CD26
Mean trimmed ADT expression per SubjectId
ns
*

a)

| Cluster | Cell Type |
|---|---|
| 0 | Predominantly CD4 |
| 1 | Predominantly CD8 |
| 2 | Mixed |
| 3 | Double Negative |
| 4 | Predominantly CD4 |
| 5 | Predominantly CD4 |
| 6 | Double Negative |
| 7 | Predominantly CD8 |
| 8 | Predominantly CD4 |

b)

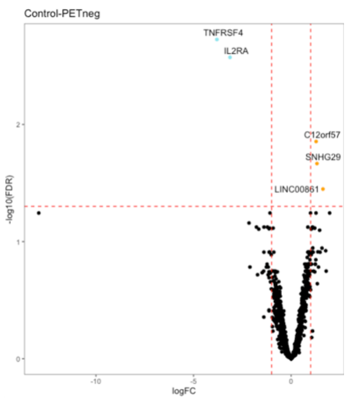


c)

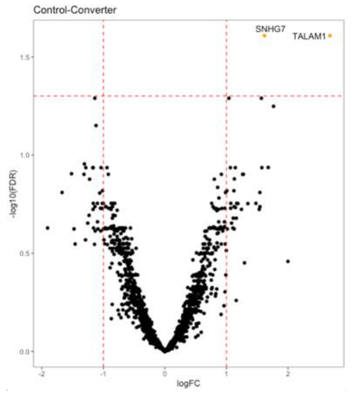


d)

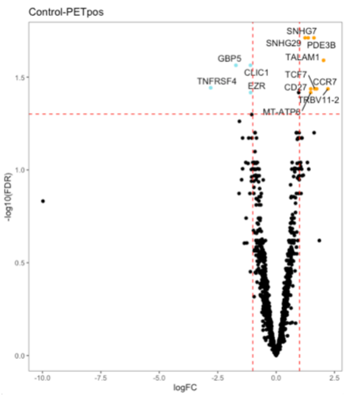


e)

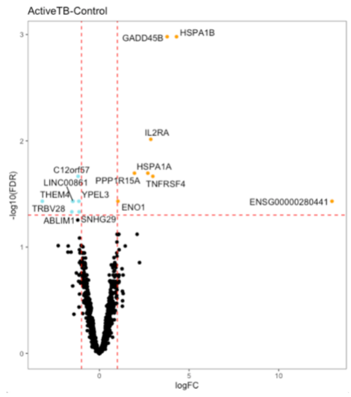


f)

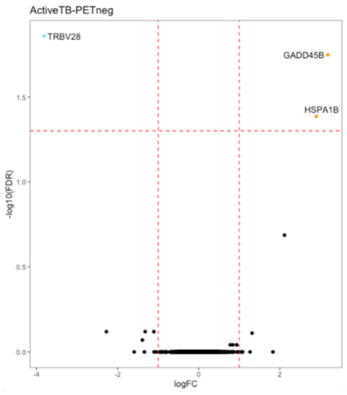


g)

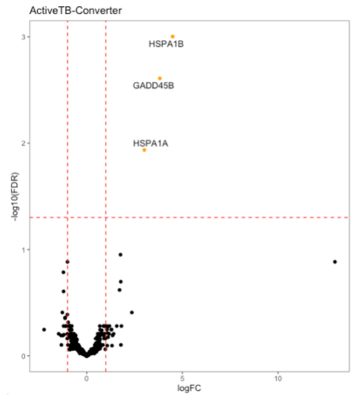


h)

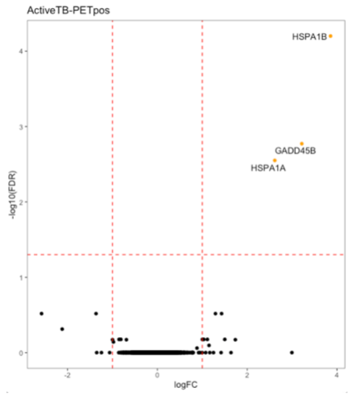

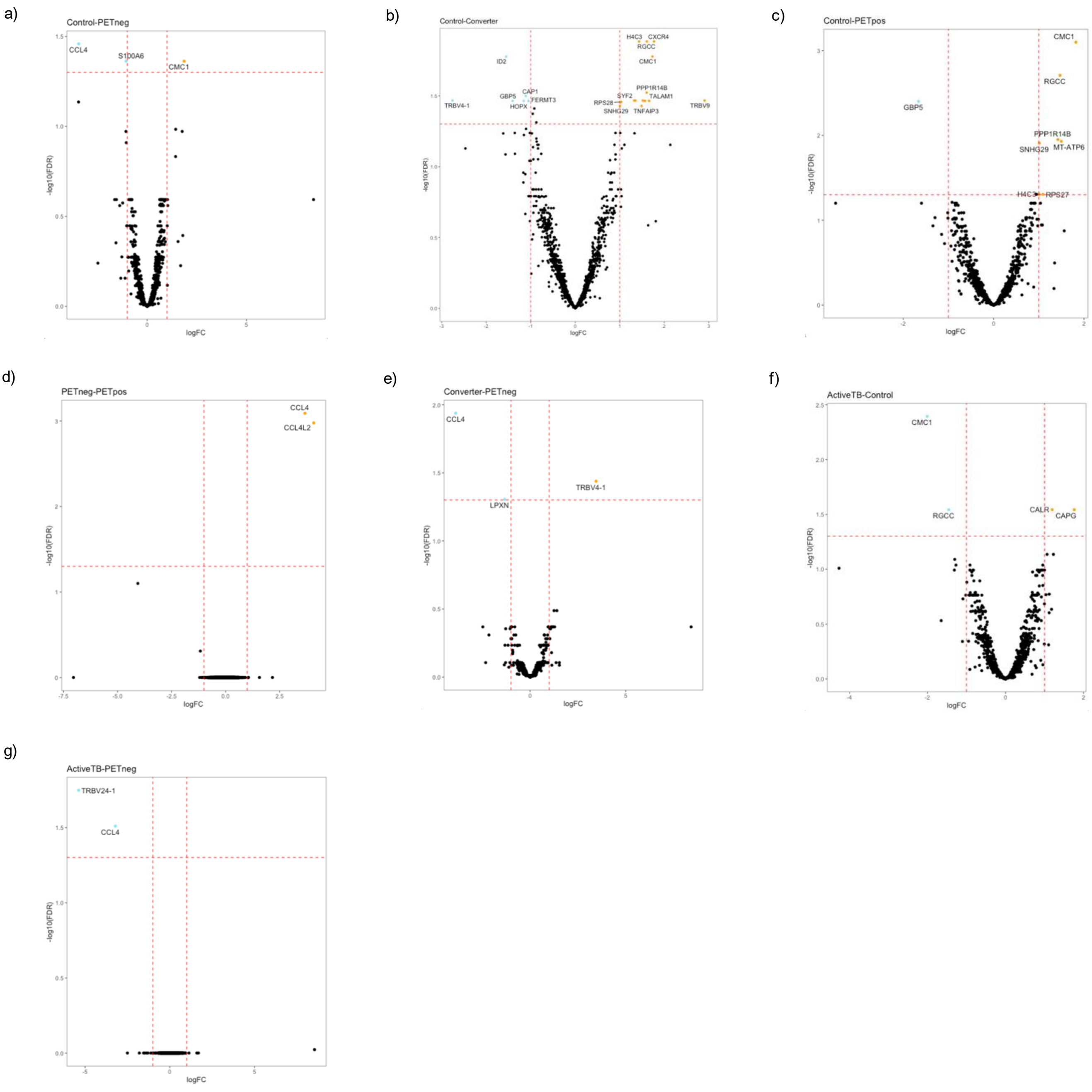
a)
Control-PETneg
CCL4
S100A6
CMC1
-log10(FDR)
logFC
b)
Control-Converter
H4C3
CXCR4
RGCC
ID2
CMC1
PPP1R14B
CAP1
GBP5
FERMT3
SYF2
TALAM1
TRBV4-1
HOPX
RPS28
SNHG29
TNFAIP3
TRBV9
-log10(FDR)
logFC
c)
Control-PETpos
CMC1
RGCC
GBP5
PPP1R14B
SNHG29
MT-ATP6
H4C3
RPS27
-log10(FDR)
logFC
d)
PETneg-PETpos
CCL4
CCL4L2
-log10(FDR)
logFC
e)
Converter-PETneg
CCL4
TRBV4-1
LPXN
-log10(FDR)
logFC
f)
ActiveTB-Control
CMC1
RGCC
CALR
CAPG
-log10(FDR)
logFC
g)
ActiveTB-PETneg
TRBV24-1
CCL4
-log10(FDR)
logFC

a)

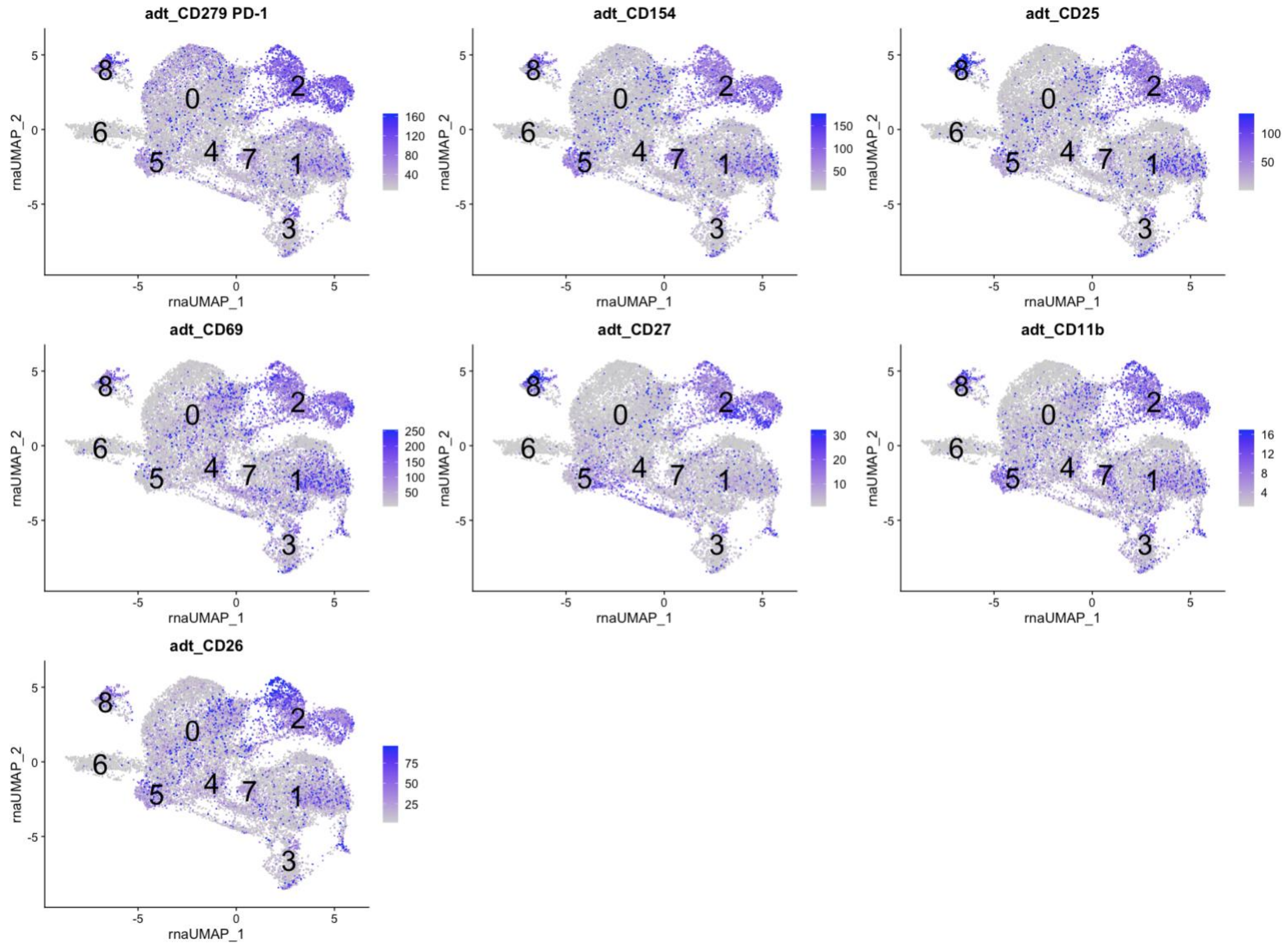
adt_CD279 PD-1
160
120
80
40
adt_CD154
150
100
50
adt_CD25
100
50
adt_CD69
250
200
150
100
50
adt_CD27
30
20
10
adt_CD11b
16
12
8
4
adt_CD26
75
50
25
rnaUMAP_1
rnaUMAP_2
0
1
2
3
4
5
6
7
8

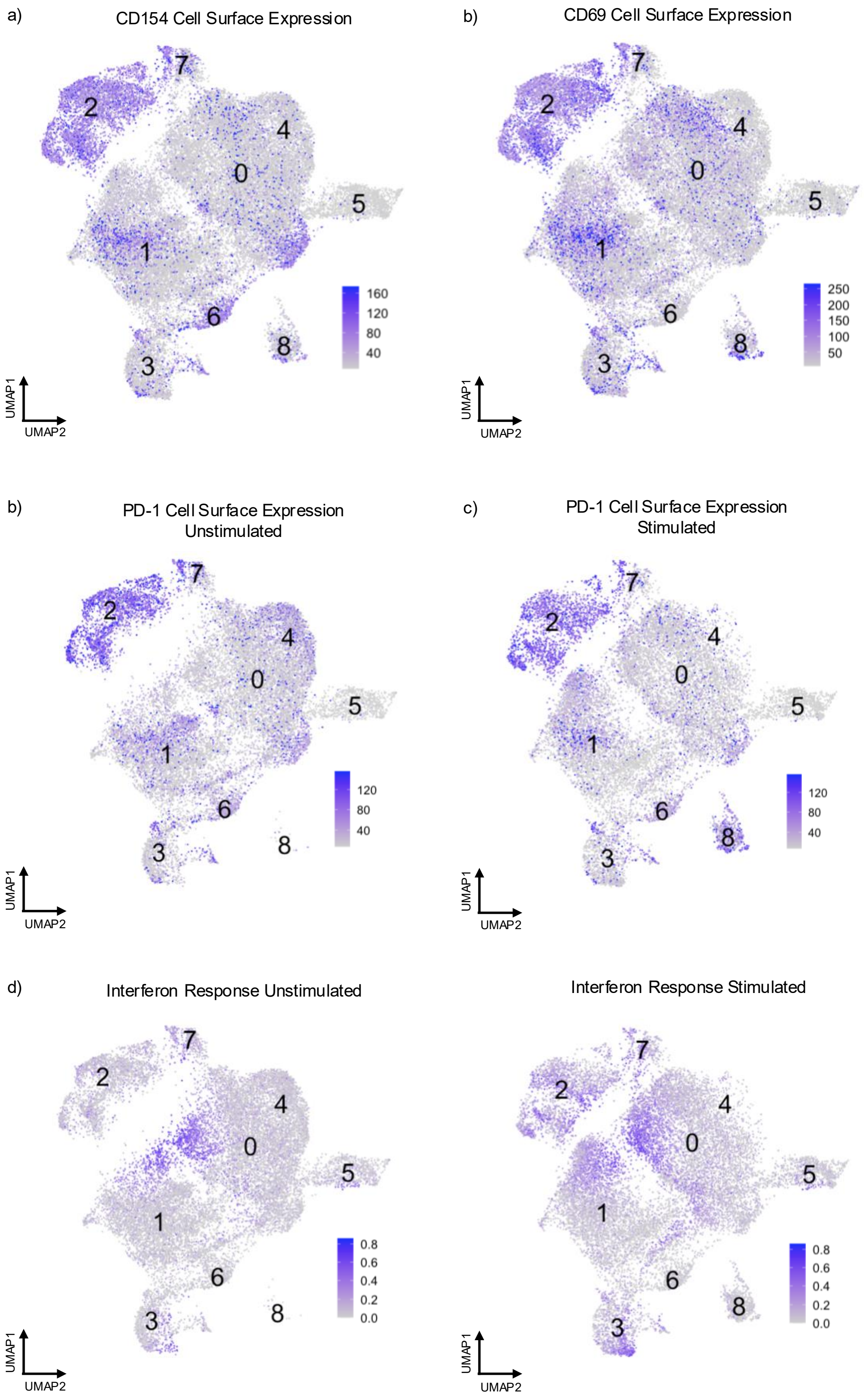
a)
CD154 Cell Surface Expression
160
120
80
40
UMAP1
UMAP2
b)
CD69 Cell Surface Expression
250
200
150
100
50
UMAP1
UMAP2
b)
PD-1 Cell Surface Expression
Unstimulated
120
80
40
UMAP1
UMAP2
c)
PD-1 Cell Surface Expression
Stimulated
120
80
40
UMAP1
UMAP2
d)
Interferon Response Unstimulated
0.8
0.6
0.4
0.2
0.0
UMAP1
UMAP2
Interferon Response Stimulated
0.8
0.6
0.4
0.2
0.0
UMAP1
UMAP2
